# Multistage and transmission-blocking tubulin targeting potent antimalarial discovered from the open access MMV Pathogen Box

**DOI:** 10.1101/2022.04.20.488926

**Authors:** Geeta Kumari, Ravi Jain, Raj Kumar Sah, Inderjeet Kalia, Manu Vashistha, Pooja Singh, Agam Prasad Singh, Kirandeep Samby, Jeremy Burrows, Shailja Singh

## Abstract

Development of resistance to current antimalarial therapies remains a significant source of concern. To address this risk, new drugs with novel targets in distinct developmental stages of Plasmodium parasites are required. In our current work, we have targeted *P. falciparum* Tubulin (*Pf*Tubulin) proteins which represent some of the potential drug targets for malaria chemotherapy. Plasmodial Microtubules play a crucial role during parasite proliferation, growth, and transmission, which render them highly desirable targets for the development of next-generation chemotherapeutics. Towards this, we have evaluated the antimalarial activity of Tubulin targeting compounds received from the Medicines for Malaria Venture (MMV) “Pathogen Box” against the human malaria parasite, *P. falciparum* (including 3D7, RKL-9 (Chloroquine resistant) and R539T (Artemisinin resistant) strains). At nanomolar concentrations, filtered out compounds exhibited pronounced multistage antimalarial effects across the parasite life cycle, including intra-erythrocytic blood stages, liver stage parasites, gametocytes and ookinetes. Concomitantly, these compounds were found to impede male gamete ex-flagellation, thus showing transmission-blocking potential of these compounds. Target mining of these potent compounds, by combining *in silico*, biochemical and biophysical assays, implicated *Pf*Tubulin as their molecular target, which may possibly act by disrupting microtubule assembly dynamics by binding at the interface of α-βTubulin-dimer. Further, promising ADME profile of the parent scaffold supported its consideration as a lead compound for further development. Thus, our work highlights the potential of targeting *Pf*Tubulin proteins in discovering and developing next-generation, multistage antimalarial agents for treating Multi-Drug Resistant (MDR) malaria parasites.

**GRAPHICAL ABSTRACT:** 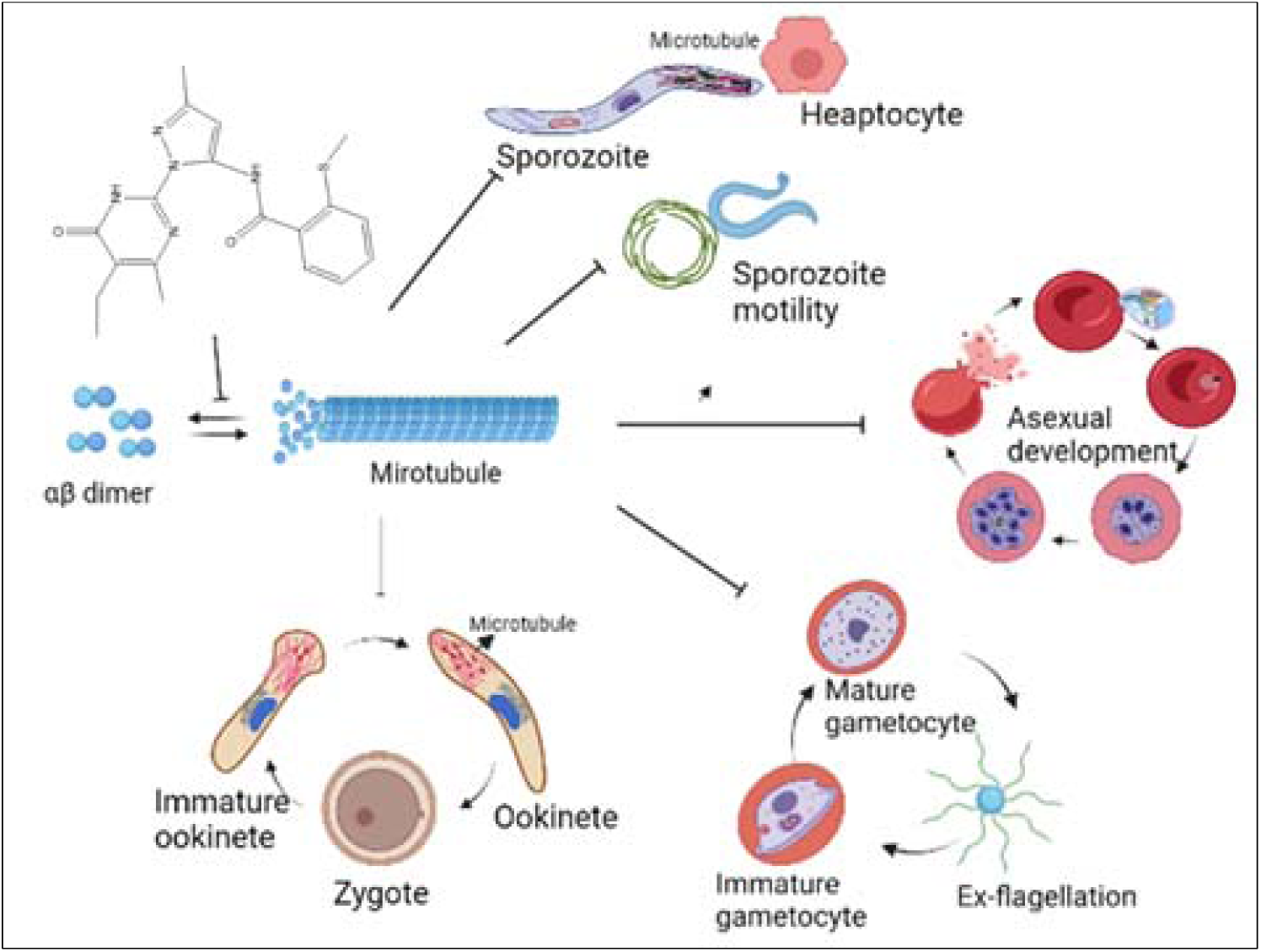

## 1. INTRODUCTION

Malaria is a global health problem, still causing 435,000 deaths each year and over 244 million cases according to 2021 WHO report [1]. Due to the lack of a fully effective vaccine, as well as the increasing threat of parasite and vector resistance, the development of new drugs that combat resistant parasites is of the highest priority.

Microtubules (MTs) are well known for their therapeutic potential in the treatment of cancer, gout, and helminth infections [2]. Likewise, tubulin and microtubules are a potential target for antiparasitic agents aimed at inhibiting growth of apicomplexan protozoan parasites [3–8].

During asexual erythrocytic development, malaria parasites undergo various morphological changes that require the cytoskeleton to be both stable and flexible [9–11]. Apart from microfilaments (actin filaments) and intermediate filaments, microtubules are another structural component of the eukaryotic cytoskeleton [12] which along with other cytoskeletal proteins such as actin, actin associated proteins, intermediate filaments and myosin are present in malaria parasites. MTs are composed from protofilaments of α-tubulin and β-tubulin, which arrange into hollow cylinders with a diameter of 25 nm [6]. MTs are polar, with one end of the microtubule crowned by α-tubulins (minus end) and the other by β-tubulins (plus end). MTs are essential for many cellular processes, including intracellular transport, chromosome segregation, establishment and maintenance of polarity, and migration [11, 12]. In *Plasmodium*, they are classified into three classes: subpellicular, spindle, and axonemal microtubules [9]. Subpellicular microtubules (sMTs) are found in all motile stages of the parasite life cycle: the sporozoites, ookinetes and merozoites [13–16].

In replicating parasites, spindle microtubules are present at the time of schizogony. This implicates not only the function of microtubules in nuclear segregation, but also in the assembly of daughter cells [17, 18]. Axonemal microtubules are exclusively found in microgametes in *Plasmodium* [19]. Similar, to other eukaryotic axonemes, the flagellum of *Plasmodium* microgametes exhibits the characteristic microtubule arrangement of 9x2 + 2 [20–23]. The flagellum is used for locomotion, to reach a macrogamete and accomplish efficient sexual replication. Given the importance of microtubules throughout development of the parasite including sporozoite, erythrocytes, gametocytes and ookinetes, targeting this would be a promising approach to inhibit malaria parasite at different stages of its life cycle. In order to be effective as antiparasitic agents, tubulin-targeting substances should be selective for the pathogen over the host target with low non-specific effects or cytotoxicity to humans.

In this study we report the antimalarial activity of tubulin targeting compounds received from the Medicines for Malaria Venture (MMV). MMV is a Product Development Partnership (PDP) in antimalarial drug research and has an extensive network of over 400 pharmaceutical, academic and endemic-country partners in more than 55 countries. Recently MM676477, a molecule from the pathogen box, was recently evaluated against *L. amazonesis* and found to increase leishmania microtubule polymerization [24]. Since tubulin and microtubules are required for the multiplication of protozoan’s parasites it can serves as a potential therapeutic target for the development of novel antiparasitic agents.

To test whether compound MMV676477 and its analogs are active against malaria parasite as well. Compounds MMV676477 and its analogs were received from MMV pathogen box. They were first analyzed for their structural similarity using *in silico* tool, Discovery Studio Visualizer which showed that 21 of them showed similarity index more than 0.8. These compounds was further investigated for their antimalarial activity against 3D7 (chloroquine and artemisinin sensitive), RKL9 (chloroquine resistant) and R539T (artemisinin resistance) cell lines of *P. falciparum* and result showed that all these compounds exhibited antimalarial activity in 150 nM to 30 µM range. Furthermore, resistance index was calculated for all the compounds, low level of cross resistance was found with 3D7-RKL9 and 3D7-R539T, suggesting these compounds inhibiting an additional target in resistant cell lines. Among the compounds tested, MMV676477, MMV1578136 and MMV1578138 showed higher antimalarial potency as well as low resistance index. Further, these compounds showed low cytotoxicity against hepatocyte mammalian cell line with selective index of more than ≥90. Profiling of MMV676477’s susceptibility to different asexual blood stages of the malaria parasite showed its peak activity against trophozoites and schizonts. The *in-silico* interaction analysis implicated Pf tubulin as a molecular target of MMV676477 which shown to bind at the α-β dimer interface. In order to validate in silico data, we have cloned and expressed Pf α-tubulin-I and β-tubulin in *E. coli* and studied its interaction with MMV676477. SPR analysis confirmed interaction between MMV676477 and Pf tubulin with Kd value of ∼ 27.7 µM. Further in vitro microtubule polymerization assay with recombinant Pf tubulin in presence of MMV676477 revealed increase level of polymerization indicating stabilization of microtubule assembly. As microtubule dynamicity play key role throughout development of malaria parasite, we next screen its effect on hepatocyte infection by sporozoite in vitro and found that this compound inhibited sporozoite infection significantly. Moreover, MMV676477 significantly inhibited gametocyte maturation, ex-flagellation of male gametocyte and ookinete maturation of *P.berghei* when tested ex vivo, demonstrating its transmission-blocking activity.

Altogether, this study gives insight into the importance of microtubule dynamicity at every stage of the *Plasmodium* life cycle. Drug that disrupts the balance between free tubulin dimers and polymerized microtubules can cause disruptions in essential processes, such as cell division, and may cause targeted cell death. In this study we demonstrated the broad antimalarial activity of MMV tubulin targeting compound against various life stages of the malaria parasite including sporozoite, erythrocytic, gametocyte and ookinete stage. Multistage activity is a desirable attribute for the next generation of antimalarials [23]. Moreover, this study identifies proteins that are effective at targeting different stages of the parasite life cycle.

## 2. MATERIAL AND METHODS

### 2.1 Structural analysis of MMV676477 & its analogs

Thirty tubulin targeting compounds, including the parent scaffold: MMV676477 and its synthetic analogs were received from Medicines for Malaria Venture (MMV) “Pathogen Box” (https://www.mmv.org/mmv-open/archived-projects/pathogen-box). MMV is an anti-malarial medication research Product Development Partnership (PDP) with a global network of over 400 pharmaceutical, academic, and endemic-country partners in over 55 countries.

Structural superimposition of MMV676477 and its analogs was done by using Discovery Studio Visualizer v20.1.0.19295, developed by Dassault Systèmes Biovia Corp. (https://www.3ds.com/products-services/biovia/), and overlay similarity was evaluated taking the parent scaffold (MMV676477) as a reference compound. Chemical structures of MMV676477 and its analogs were drawn by using ChemDraw Ultra v12.0.2.1076, one of the CambridgeSoft products for producing a nearly unlimited variety of biological and chemical drawings.

### 2.2 *Pf*αI-β tubulin architecture

Comparative or homology modeling was employed to model 3D structure of *Pf*αI-β tubulin heterodimer, by following the protocol as described previously by our group [25],[26]. To accomplish this, amino acid sequences of *Pf*αI- and *Pf*β-tubulins (PlasmoDB IDs: PF3D7_0903700 and PF3D7_1008700, respectively) were retrieved from PlasmoDB database [27]. To search for a suitable template for homology modeling, BLASTp search was performed using amino acid sequences of *Pf*αI- and *Pf*β-tubulins as query sequences, against Protein Data Bank (PDB) database [28],[29]. Amino acid sequence identities among tubulin orthologs from *P. falciparum* strain 3D7 and *Bos taurrus* were found to be 84.1% (α-tubulin) and 88.8% (β-tubulin), thus rendering X-Ray diffraction based structural model of *Bt*α-β tubulin heterodimer (PDB ID: 1JFF; available at resolution: 3.5 Å) as a suitable template to model 3D structure of *Pf*αI-β tubulin [30]. Homology modeling was done by using Modeler v9.17 software, a tool used for homology or comparative modeling of protein three-dimensional structures by satisfaction of spatial restraints [31]. Generated structural model was further subjected to structural refinement by using ModRefiner, which is an algorithm for atomic-level, high-resolution protein structure refinement [32]. Reliability of the refined structural model of *Pf*αI-β tubulin was assessed by examining backbone dihedral angles: phi (Ø) and psi (Ψ) of the amino acid residues lying in the energetically favorable regions of Ramachandran space [33]. This was done by using PROCHECK v.3.5, a tool which checks the stereochemical quality of a protein structure, producing a number of PostScript plots analyzing its overall and residue-by-residue geometry [34]. Percent quality assessment of the protein structure was evaluated by utilizing four sorts of occupancies: ‘*core*’, ‘*additionally allowed*’, ‘*generously allowed*’ and ‘*disallowed regions*’. The refined 3D structural model of *Pf*αI-β Tubulin heterodimer, thus generated, was subsequently used for *in silico* interaction analysis with MMV676477 and its analogs.

### 2.3 *In silico* interaction analysis of MMV676477 and its analogs with *Pf*αI-β tubulin

Energy-minimized 3D structural models of MMV676477 and its analogs were generated by using Chem3D Pro 12.0 (https://perkinelmerinformatics.com/products/research/chemdraw/), as described previously by our group [26]. Molecular docking studies were performed by using Autodock Vina Tools 1.5.6 to rationalize the inhibitory activities of MMV676477 and its analogs against *Pf*αI-β tubulin [35]. We ensured that the whole tubulin heterodimer was covered while constructing the virtual grid for docking. For this, a virtual 3D grid of 64×70×90, with x, y, z coordinates of the center of energy: 227.06, 386.225 and 192.023, respectively, was constructed using Autogrid module of AutoDock Tools. Top scoring docked conformations of MMV676477 and its analogs were selected based on their most negative free binding energies and visualized for polar contacts (if any) with the amino acid residues of *Pf*αI-β tubulin by using PyMOL Molecular Graphics System [36].

A scatter graph was used to show the association (if any) between the two variables: *in silico* binding energies, *ΔG_bind_* (kCal/mol) of MMV676477 and its analogs with *Pf*αI-β tubulin heterodimer, and overlay similarity among the compounds (taking MMV676477 as the reference scaffold), by using Graph Pad Prism 8.0.1, GraphPad Software, San Diego, California USA (www.graphpad.com).

### 2.4 SPR analysis of binding affinity between *Pf*αI- and *Pf*β-tubulins

Cloning, over-expression and purification of *Pf*αI- and *Pf*β-tubulins (PlasmoDB IDs: PF3D7_0903700 and PF3D7_1008700, respectively) was done, as described previously by our group [37]. To determine binding strength of *Pf*αI- and *Pf*β-tubulin *in vitro*, real-time biomolecular interaction analysis was carried out with SPR at physiologically relevant concentrations, by using AutoLab Esprit SPR (at Advanced Instrumentation Research Facility, Jawaharlal Nehru University, New Delhi, India), and affinity constant, K_D_ was determined at Room Temperature (RT, 298K). SPR analysis was performed by following previously described protocols [38–42]. Briefly, interaction kinetics was studied by injecting *Pf*β-tubulin (in HEPES-NaCl buffer, pH 7.5) at different concentrations: 25, 50, 100, 200, and 400 nM over the *Pf*αI-tubulin (in the same buffer) immobilized (5.2 ng per mm^2^) immobilized SPR sensor chip surface (self-assembled monolayer of 11-Mercapto-Undecanoic Acid, MUA on gold surface) by the mechanism of “covalent amine coupling”, with association and dissociation time of 500 s and 300 s, respectively, followed by comparing their respective binding affinities at RT. A control flow cell was activated and blocked in a similar manner with injection of buffer equivalent concentrations of DMSO to allow for reference subtraction. HEPES-NaCl buffer, pH 7.5 was used both as immobilization and binding solutions. The sensor chip surface was regenerated with 50 mM NaOH. Data were fit to the two-state conformational change model using AutoLab SPR Kinetic Evaluation software provided with the instrument. K_D_ values were calculated using the Integrated Rate Law (IRL) equation.

### 2.5 SPR analysis of binding affinity between MMV676477 & its analogs, and *Pf*-tubulins

To determine binding strength of MMV676477 & its analogs (n = 30) *in vitro*, real-time biomolecular interaction analysis was carried out with SPR, as described above. Briefly, all compounds were injected individually at different concentrations: 75, 50, 25, 6.25, 1.56 and 0.78 μM over the *Pf*αI- and *Pf*β-tubulin immobilized SPR sensor chip surfaces, with association and dissociation time of 300 s and 200 s, respectively, followed by comparing their respective binding affinities at RT. A control flow cell was activated and blocked in a similar manner with injection of equivalent concentrations of DMSO to allow for reference subtraction. HEPES-NaCl buffer was used both as immobilization and binding solutions. Data were fit to the two-state conformational change model using AutoLab SPR Kinetic Evaluation software provided with the instrument. K_D_ values were calculated using the Integrated Rate Law (IRL) equation.

### 2.6 Microtubule assembly assay

*Pf*-tubulin polymerization was carried out, as described previously, in 200 *μ*l of a ice-cold re-assembly buffer (100 mM PIPES, pH 6.9, 10 % glycerol, 2 mM MgCl_2_.H_2_O, 0.5 mM EGTA), containing 1 mM GTP, different equimolar concentrations (0.5, 1.0, 2.5, 5, 10, 20 and 40 *μ*M) of *Pf*αI- and *Pf*β-tubulins [43–47]. The reaction mixture was prepared in clear, flat bottom 96-welled plates. The time-course of tubulin polymerization was followed over 30 min at 37°C, by monitoring a change in turbidity at 350 nm using a Varioskan™ LUX multimode microplate reader (Thermo Scientific™) equipped with an incubator for temperature control.

To investigate the effect of MMV676477 and its analogs on *Pf*-tubulin polymerization, 200 μl of the tubulin reaction mixture (containing 10 μM of *Pf*αI- and *Pf*β-tubulins), was added to the 96-welled plate along with 10 μM of each compound, and the time-course of tubulin polymerization was followed over 18 min.

### 2.7 GTPase Assay

GTPase activity contributed by *Pf*β-tubulin in the microtubule polymerization assay reactions was evaluated by measuring the amount of P_i_ produced in the reaction mixtures (at 18 min.) by using Malachite Green Phosphate Assay Kit (BioAssay Systems, POMG-25H), that relies on estimation of green colored complex formed between Malachite Green, molybdate and free orthophosphate (PO_4_^3−^) released during a particular GTPase or ATPase reaction. Protocol was followed as described previously [48]. Green color formation in the reaction mixtures was measured after 20 minutes using a Varioskan™ LUX multimode microplate reader (Thermo Scientific™).

### 2.8 Parasite culturing

*In vitro* Culture of *Plasmodium falciparum* 3D7 strain was cultured using O+ human erythrocytes, under mixed gas environment (5% O2, 5% CO2, and 90% N2) as described previously [49]. The culture media was composed of RPMI 1640 (Invitrogen, Carlsbad, CA, United States), supplemented with 50 mg/L hypoxanthine (Sigma-Aldrich, St. Louis, MO, United States), 2 g/L sodium bicarbonate (Sigma-Aldrich, St. Louis, MO, United States), and 5 g/L Albumax I (Gibco, Grand Island, NY, United States). Human erythrocytes (O+) as well as the serum of healthy volunteers were procured from the blood bank and used only for research purposes. Synchronous development of the erythrocytic stages of a human malaria parasite, *P. falciparum*, in culture was accomplished by suspending cultured parasites in 5% D-sorbitol and subsequent reintroduction into culture [50].

### 2.9 *In vitro* antimalarial activity of MMV compounds against human malaria parasite

Three different strains of *P. falciparum*, RKL-9 (chloroquine-resistant), 3D7 (chloroquine-sensitive) and R539T (artemisinin-resistant) were used for the chemosensitivity tests. For this the compounds were dissolved in DMSO and then diluted with medium to achieve the required concentrations (final DMSO concentration <1%, which is non-toxic to the parasite). The drugs were placed in 96-wells flat-bottom microplates in duplicate at different concentration (20, 10, 5, 2.5, 1.25, 0.625, 0.312, 0.156µM). Sorbitol synchronized cultures with 0.8-1% parasitemia and 2% hematocrit were aliquoted into the plates and incubated for 72 h in a final volume of 100 µL/well. Chloroquine and artesunate was used as a reference compound. Parasite growth was determined with SYBR Green I based fluorescence assay. Briefly, after 72 h of incubation culture was lysed by freeze-thaw. Followed by addition of 100 µL of lysis buffer (20 mM Tris/HCl, 5 mM EDTA, 0.16% (w/v) saponin, 1.6% (v/v) Triton X) containing 1× SYBR Green I ((Thermo Fisher Scientific, Waltham, Massachusetts, US)). Plates were incubated in the dark at room temperature (RT) for 3-4 h. *P. falciparum* proliferation was assessed by measuring the fluorescence using a Varioskan™ LUX multimode microplate reader (Thermo Scientific™) with an excitation and emission of 485 nm & 530 nm respectively. IC50 values were determined via non-linear regression analysis using GraphPad prism 8.0 software. The results were expressed as the percent inhibition compared to the untreated controls, calculated with the following formula: 100× ([OD of Untreated sample – blank]-[OD– blank]/ [OD – blank]). As blank, uninfected RBC were used. IC50, which is the dose required to cause 50% inhibition of parasite viability, was determined by extrapolation

### 2.10 Stage specific activity of MMV compound against asexual stage of *Plasmodium falciparum*

Standard asexual blood stage susceptibility results were collected by exposing asynchronous 3D7 parasite cultures to 7 different concentrations (10, 5, 2.5, 1.25, 0.625, 0.312, 0.156 µM) plus no-compound controls for 72 h. To determine the specific asexual blood stage at which the compounds are active, schizonts were percoll-purified **(54)** following consecutive synchronization with 5% sorbitol [51]. As culture was allowed to re-invade and synchronize with sorbitol again These parasites were then plated in 96-well plates at 2% hematocrit, 1% parasitemia and exposed to different concentration of MMV676477 at early rings (0-8 h), early trophozoites (16-24 h), and early schizonts (32-40 h). Incubation times were adjusted to the 40-h asexual blood stage cycle of the 3D7 parasite line. Compounds were removed through five rounds of washing after 10 h of incubation and kept again in humidified chamber. Growth inhibition was assessed at the 60-h time point at which parasites had expanded, reinvaded new RBCs, and developed into the trophozoite stage that allows straight-forward quantification by SYBR-Green I based fluorescence assay. IC50 values were determined from growth inhibition data using nonlinear regression analysis (Prism 8, GraphPad). All asexual blood stage assays were repeated on at least three independent occasions with two technical replicates.

### 2.11 Cellular thermal shift analysis of binding between MMV compound and its target molecule

The binding of compound MMV676477 with Pf-tubulin was confirmed by cellular thermal shift assay (CETSA) which is a method for examining ligand binding with the target protein in cell lysate [52, 53]. Samples for CETSA studies were prepared using highly synchronous schizont stage *P. falciparum* 3D7 parasites. Culture at 10% parasitemia was pelleted down and lysed with 0.03% saponin (Sigma-Aldrich) in ice-cold 1XPBS and incubated on ice for 10 min. Parasites were washed 3 times in ice-cold PBS and final pellet was resuspended in 10 volumes of lysis buffer (1X PBS, 1% Triton X-100, 1% NP-40,) and lysed by three freezing-thawing cycles. The lysates were cleared by centrifugation at 16,000 g for 30 min and the supernatant fraction was transferred to fresh tube.

The parasite lysate was then subjected to drug treatment for 30 min and as a control, lysate were left untreated. After 30 min of incubation the fraction of lysate were transferred to tubes and heated at respective temperatures for 3 min in C1000 Touch PCR thermal cycler (Bio-Rad), followed by 3 min incubation at 4°C. The post-heating lysates were centrifuged at 13,000 g for 30 min at 4°C. Soluble proteins were resolved on SDS-PAGE gel and electrophoretically transferred to Immun-Blot® PVDF Membrane (Bio-Rad) using Trans-Blot® SD Semi-Dry Transfer Cell (Bio-Rad). Blot were blocked with 5% non-fat milk in 1X TBST for 1 h and probed with anti-alpha tubulin (produced in-house; 1:2000) for 1 h, followed by washing with 1X PBST thrice. After washing blot were incubated with HRP-conjugated **anti-rabbit IgG Ab** (1: 10,000) for 1 h and washed again with 1X PBST thrice. Blots were developed with ECL Western Blotting Substrate (Thermo Fisher Scientific) and analyzed using ChemiDoc Gel Imaging System and Image Lab Software (Bio-Rad).

### 2.12 Immunofluorescence assay

Immunofluorescence assay (IFA) was performed to see the effect of MMV676477 on dynamicity of microtubule structure in Plasmodium parasite. Late stage schizont were treated with MMV676477 at their IC50 concentration for 6 h. After incubation, thin smear of schizont stage were made on glass slide, air dried, and fixed with methanol (ice cold, 100%) for 30 min at −20°C. slides were then blocked for 1 h at room temperature (RT) in 3% bovine serum albumin (BSA) made in 1% PBS. After blocking, slides were then first incubated with anti-alpha tubulin (produced in-house; 1:500) for 1 h at RT. Slides were then washed with 1XPBST (0.05% Tween-20) twice, followed by probing with Alexa Fluor 488 conjugated goat anti-rabbit IgG (1:500 dilution; Molecular Probes, United States) at RT for 1 h. Slides were then washed again and mounted with Pro Long Gold antifade reagent (Invitrogen, Carlsbad, CA, United States) with coverslips and viewed under Nikon A1R confocal microscope at 100X objective.

To see the effect of MMV676477 on microtubule dynamicity at sporozoite stage. Isolated sporozoites were treated with MMV676477 at 3 µM concentration for 1 h. For control sporozoite were left untreated. After treatment smear were made on glass slide and processed same as above for immunofluorescence analysis.

### 2.13 Sporozoite

*Anopheles stephensi* mosquitoes infected with *P. berghei* were anesthetized, and their salivary gland were isolated by dissection, to release the sporozoite, salivary glands were placed into complete media (DMEM (Gibco™) with 1% FBS (Gibco™) and 1X Pencillin-Streptomycin antibiotic (Gibco™)) and mechanically homogenized [6]. The homogenized solution containing sporozoite was briefly centrifuged at 1000 rpm to separate sporozoite from tissue. Supernatant fraction containing sporozoite were then resuspended in activation media (DMEM with 10% FBS and 1X pencillin/streptomycin antibiotic). Prior incubation with hepatocyte, sporozoite were counted using hemocytometer.

To study the effect of MMV67677, MMV1578136 and MMV1578138 on hepatocyte infection by sporozoite. Hepatocyte cell line HEPG2 were seeded in 12-well plate at a density of 2 × 10^5^ cells/well 20-26 h prior to the infection. Isolated sporozoite were then added to each well at 0.2 × 10^5^ density along with the MMV compound at 3 µM concentration for 3 h. After 3 h of incubation, culture was washed and incubated with fresh media containing MMV compound and further incubated for 48 h.

To estimate the sporozoite infection in hepatocyte, total RNA was extracted using TRIzol RNA Isolation Reagents (Invitrogen). Isolated total RNA was then treated with DNase I (Invitrogen^TM^) to remove DNA contamination. Purified RNA were then subjected to reverse transcription using iScript^TM^ cDNA synthesis kit (Bio-Rad). The resulting cDNA was diluted 1:10 with nuclease-free water. Real-time PCR analysis was performed using the SYBR Green qPCR Master Mix (Applied Biosystem). Briefly, the PCR reaction consisted of 4 µl of Brilliant II SYBR Green qPCR Master Mix, 10 µM of forward and reverse primers and 2 µl of diluted cDNA to a total volume of 25 µl in a MicroAmp Fast Optical 96-Well Reaction Plate (Applied Biosystems) covered with MicroAmp Optical Adhesive Film. The reaction plate was then placed in a 7,500 Fast Real-Time PCR System (Applied Biosystems) to quantify parasite 18S rRNA using the default thermal cycling conditions: 95 °C, 10 minutes followed by 40 cycles of 94 °C, 30 seconds; 55 °C, 1 minute; and 72 °C, 30 seconds. The housekeeping Beta actin gene of human was used as an endogenous control (normalizer) for every run. Results were analyzed using the Applied Biosystems standard PCR calculation system once each run was completed.

### 2.14 Sporozoite gliding motility assay

To test the effect of MMV compounds on sporozoite motility, sporozoite gliding motility assay was performed. Sporozoite isolated from salivary gland of infected mosquitoes were resuspended in activation medium. To see the effect of MMV compound on sporozoite motility, compound MMMV676477, MMV1578136, MMV1578138 was added promptly to the sporozoites to give a final concentration of 3 µM and mixed by gentle pipetting. For the control sporozoite were left untreated. Sporozoites were then allowed to glide on glass surface for 30’ at 37°C. After incubation slides were fixed with methanol and blocked with 1% bovine serum albumin (BSA) in 1× phosphate buffered saline (PBS) at RT for 1 h. Slides were probed with monoclonal CSP antibody (1:1000 in 1% BSA) for 1 h at room temperature followed by staining with anti-mouse Alexa Fluor® 488 conjugate for 1 h at RT. After incubation slides were washed thrice to remove unbound antibodies and mounted with DAPI-antifade (Invitrogen, Carlsbad, CA, USA). Illuminated anti-CSP trails were imaged by Carl Zeiss Microscopy, LLC.

### 2.15 Animal handling

The studies were conducted in compliance with guidelines established by the Institutional Animal Ethics Committee (IEAC) of Jawaharlal Nehru University (JNU), Delhi and Committee for Control and Supervision of Experiments on Animals (CPCSEA). IAEC-JNU’s animal ethics committee approved the regulations for the use of animals in research and in experimental research under strict adherence to ethical standards. In the experiment, mice were housed under standard conditions of food, temperature (25 ± 3 °C), relative humidity (55 ± 10%) and illumination (12 h light/dark cycles) obtained from the Central Laboratory Animal Resources, Jawaharlal Nehru University, Delhi.

### 2.16 *Plasmodium berghei* ANKA (PbA) infection in BALB/c mice

The experimental BALB/c mice were infected with the chloroquine-sensitive Plasmodium berghie ANKA strain (PbA). PbA-infected mice served as the source of donors. Infected Donor mice having parasitemia 20-30% were sacrificed, and blood was collected into heparinized tubes by cardiac puncture. Afterwards, the blood was diluted with 0.9% normal saline solution, and infection was induced via IP injection32 by injecting 0.2% of the diluted blood containing 1X10^8^ parasitized erythrocytes. Microscopically, parasitemia was measured daily by examining thin blood smears stained with Giemsa, and was calculated using the following formula, % parasitemia = Number of parasitized erythrocytes X 100/ Total number of parasites in the blood

### 2.17 Effect of MMV compound on *P. berghei* exflagellation assay ex-vivo

To study the effect of these compound on *P. berghei* male gametocyte ex-flagellation, mice were treated with phenylhydrazine 30 mg/kg through intraperitoneal (I.P) route to induce erythropoiesis [54]. Three days post-treatment mice were infected with 1.0 × 10^7^ *P.berghei* ANKA parasites, collected from donor mice mentioned above through I.P . Gametocytemia typically peaks three days after infection. Three days after, 120 μL of infected blood was obtained from mice and mixed with complete RPMI. Infected RBCs were then treated with 3μM of MMV676477, MMV1578136 and MMV1578138 respectively, for control infected RBCs were left untreated. Samples were kept at 37°C and incubated for 1h. After incubation drug treated blood samples were washed and mixed immediately with 200 μL of ex-flagellation medium (RPMI1640 containing 25 mM HEPES, 20% FBS, 10 mM sodium bicarbonate and 50 mM xanthurenic acid at pH 8.0) and kept at 20°C for 15 min [55, 56]. Ex-flagellation centres were then counted in 10-12 field using 63X objective of Carl Zeiss Microscopy, LLC.

### 2.18 *P. berghei* ookinete drug assay

To evaluate the effect of these compound on ookinete maturation same proportional of *P.berghei* infected RBCs were obtained from phenyl hydrazine treated mice. Infected RBCs were then mixed with ookinete media (RPMI1640 containing 25 mM HEPES, 20% FBS, 10 mM sodium bicarbonate (Sigma-Aldrich) and 50 mM xanthurenic acid (Sigma-Aldrich) at pH 8.4), maintaining 10% hematocrit along with each of the MMV compound at 3 μM concentration (MMV676477, MMV1578136 and MMV1578138) for 21-24 h at 21°C [57]. ookinete development was followed by Giemsa-stained smears.

### 2.19 Cytotoxicity evaluation of MMV compounds using tetrazolium (MTT) assays against mammalian cell line

The cytotoxicity of the MMV676477 and their analogues was estimated against Human hepatocyte cell line (HePG2) using colorimetric based MTT assay. HePG2 cells were maintained in culture flasks supplemented with DMEM, 10% (v/v) FBS, 3.7 g/l NaHCO3 (Sigma) along with 1X Penicillin-Streptomycin antibiotic and grown at 37 °C in a CO2 incubator (5% CO2). Cells were inoculated into flat-bottomed clear 96-well microplate at a density of 20000 cells in 200 µL culture medium and incubated for 24 h. Next, cells were treated for 48 h with different concentrations of given compounds (50 µM, 30 µM, 20 µM and 10 µM) in triplicate and kept at 37 °C and 5% CO2. Cells without any drug treatment were used as positive control. The cytotoxicity assay was conducted after the incubation time, cells were washed with PBS and 20 µL of 3-[4,5-dimethylthiazol-2-yl]-2,5-diphenyltetrazolium bromide ((MTT) Sigma-Aldrich) solution (5 mg/ml in 1XPBS) was added to each well, incubation were continued for an additional 3 h at 37 °C and 5% CO2. After incubation, medium was removed and 100 µL of DMSO was added to each of the well. Plate was gently rotated on an orbital shaker for 10 min to completely dissolve the precipitation. The absorbance at 570 nm was measured using Varioskan™ LUX multimode microplate reader (Thermo Scientific™) to calculate the cell viability.

### 2.20 ADME profiling of MMV676477

The examination of a chemical lead’s pharmacological properties is critical for early selection or identification. As a result, MMV676477 was chosen to determine many crucial in vitro Absorption, Distribution, Metabolism, and Excretion (ADME) parameters, including lipophilicity, solubility, hepatic microsomal stability, plasma stability, and whether the tested chemical binds to plasma protein. By evaluating these factors, we can have a better understanding of a compound’s bioavailability in the body. Another parameter is the profiling of hERGs (human ether-a-go-go related genes), which are involved in cardiac potential repolarization. All these parameters were validated using MMV676477 in accordance with the established protocols [58].

## 3. RESULTS

### 3.1 Structural similarity among MMV676477 and its analogs

Structural superimposition of MMV676477 and its analogs, using BIOVIA Discovery studio Visualizer, depicted that the majority of the analogs have a strong structural similarity with the parent scaffold, MMV676477 **(Figure 1A (i))**. This has been graphically demonstrated by plotting similarity overlays with MMV676477 (taken as a reference) on the Y-axis. Accordingly, 21 analogs were found to have an overlay similarity index more than 0.8 **(Figure 1A (ii))**. To investigate if these structural similarities or differences among MMV676477 and its analogs are reflected in their interaction behavior with *Pf*αI and *Pf*β tubulins, both *in silico* and *in vitro* interaction analysis were performed.

**Figure 1:**
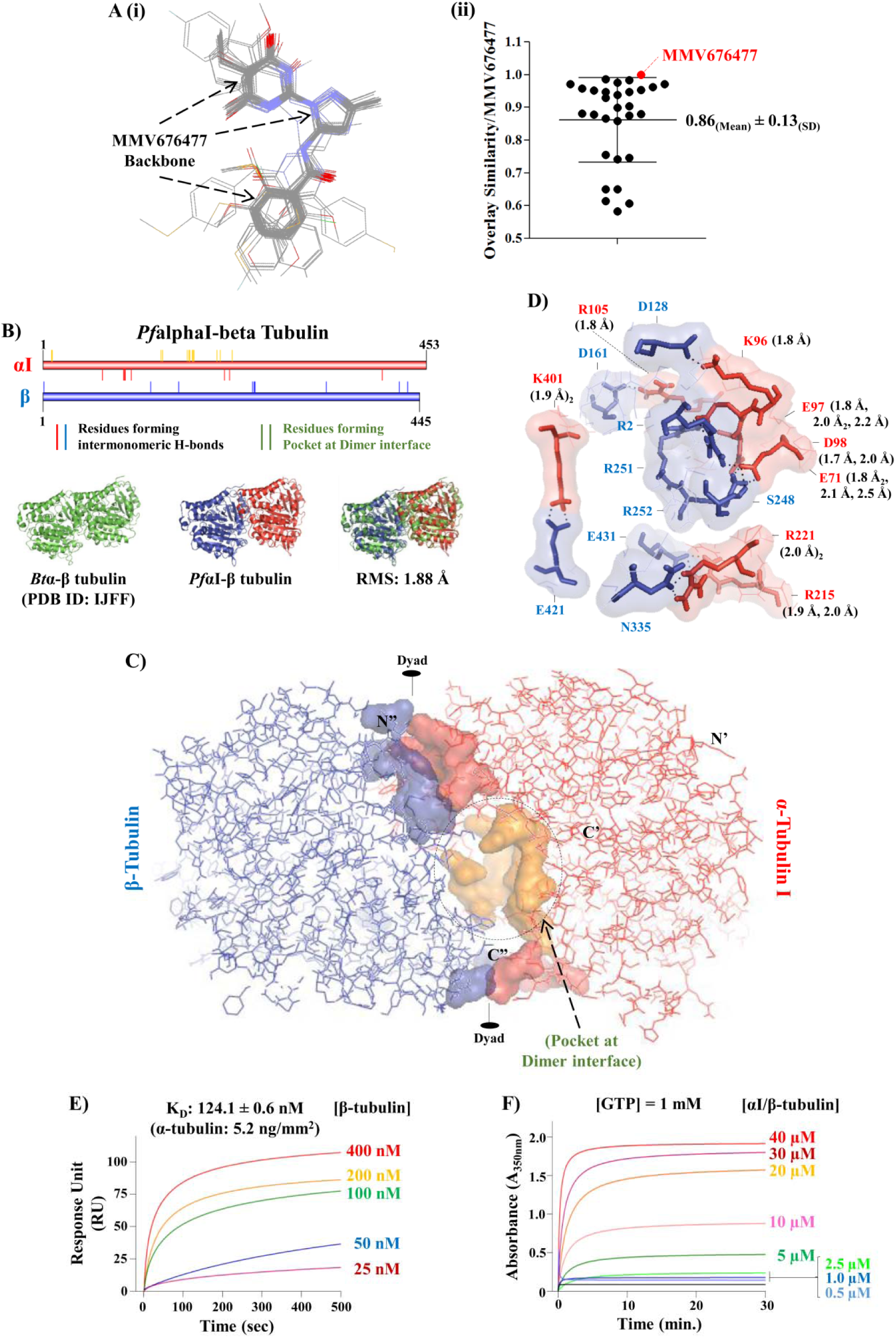
**(a) Structural similarity among MMV676477 and its analogs.** Structural superimposition of MMV676477 and its analogs depicted that the majority of the analogs have a strong structural similarity with the parent scaffold, MMV676477 **(i)**. Graphical representation of overlay similarity with MMV676477 on the Y-axis. 21 analogs were found to have an overlay similarity index more than 0.8 **(ii)**. **(b-d) Structural model of *Pf*αI-β tubulin heterodimer.** Optimal rigid-body superimposition of *Bt*αtgβ tubulin with the generated structural model of *Pf*αfiβ tubulin is shown. Overall RMSD value of the C-alpha atomic co-ordinates was found to be 1.88 Å, suggesting a reliable 3D structural model of *Pf*αfrβ tubulin. Inter-monomeric H-bond interactions are also shown. **(e) SPR-based interaction analysis of *Pf*αI and *Pf*β-tubulins.** Concentration dependent real-time SPR sensorgram to quantify the interaction between *Pf*αI and *Pf*β-tubulins is shown. With increase in mass concentration of *Pf*β-tubulin, gradual increase in sensor signal was observed, which linearly correlated with corresponding change in refractive index of the medium immediately adjacent to the SPR sensing surface. *Pf*αI-tubulin showed binding affinity for *Pf*β-tubulin, with a K_D_ value of 124.1 ± 0.6 nM. **(f) *In vitro Pf*microtubule assembly assay.** Induction of *Pf*αI/*Pf*β-tubulin assembly into microtubules, suggesting that Plasmodial *Pf*αI and *Pf*β-tubulins behave similar to the tubulins from other biological systems.

### 3.2 Structural model of *Pf*αI-β tubulin heterodimer

Owing to the high sequence similarity between tubulin orthologs from *P. falciparum* strain 3D7 and *Bos taurrus*, the X-Ray diffraction based structural model of *Bt*α-β tubulin heterodimer (PDB ID: 1JFF) was used as a template to generate three-dimensional co-ordinates of *Pf*αI-β tubulin by homology modeling, and thus, inter-monomeric H-bond interactions were elucidated (**Figure 1B, 1C and 1D**). After optimal rigid-body superimposition of *Bt*αtgβ tubulin with the generated structural model of *Pf*αfiβ tubulin, overall Root-Mean-Square Deviation (RMSD) value of the C-alpha atomic co-ordinates was found to be 1.88 Å, suggesting a reliable 3D structural model of *Pf*αfrβ tubulin (**Figure 1B**). The assessment of stereochemical quality and accuracy of the generated structural model displayed 88.9% of amino acid residues lying in the most favored (*core*) regions, with 10.5%, 0.4%, and 0.1% residues in *additional allowed*, *generously allowed* and *disallowed regions* of the Ramachandran plot, respectively, indicating that the backbone dihedral angles: ɸ and Ψ of the generated structural model of *Pf*αfrβ tubulin were reasonably accurate. RMSD value and Ramachandran plot characteristics confirmed reliability of the homology-based structural model of *Pf*αI/β tubulin to be taken further for *in silico* interaction analysis.

### 3.3 SPR-based interaction analysis of *Pf*αI and *Pf*β-tubulins

Cloning, over-expression and purification of *Pf*αI- and *Pf*β-tubulins (PlasmoDB IDs: PF3D7_0903700 and PF3D7_1008700, respectively) was done, as described previously by our group [37]. To quantitate the interaction between *Pf*αI and *Pf*β-tubulins, SPR analysis was performed by using AutoLab Esprit SPR. *Pf*αI-tubulin was immobilized at an average density of 5.2 ng per 1 mm^2^ of the SPR sensor chip surface. Once immobilized, *Pf*αI-tubulin demonstrated good stability throughout the experiment. Interaction analysis was done by injecting serial dilutions of *Pf*β-tubulin, ranging from 25 nM to 400 nM, over the *Pf*αI-tubulin-immobilized sensor chip surface, followed by comparing their respective kinetics & binding affinities at RT. With an increase in mass concentration of *Pf*β-tubulin, gradual increase in sensor signal was observed, which linearly correlated with corresponding change in refractive index of the medium immediately adjacent to the SPR sensing surface. *Pf*αI-tubulin showed binding affinity for *Pf*β-tubulin, with a K_D_ value of 124.1 ± 0.6 nM. Concentration dependent real-time sensorgram along with K_D_ value of the interaction is shown in **Figure 1E**.

### 3.4 *In vitro Pf*microtubule assembly assay

Next, we measured the polymerization time-course of *Pf*microtubules in an assembly buffer containing 1 mM GTP and different equimolar concentrations (0.5, 1.0, 2.5, 5, 10, 20 and 40 *μ*M) of *Pf*αI- and *Pf*β-tubulins. The reaction mixture was incubated at 37°C, and the turbidity changes over 30 min were determined at 350 nm using a spectrophotometer. We observed potent induction of *Pf*αI/*Pf*β-tubulin assembly into microtubules, suggesting that Plasmodial *Pf*αI and *Pf*β-tubulins behave similar to the tubulins from other biological systems **Figure 1F**. For the subsequent microtubule assembly based experiments, the concentration of *Pf*αI and *Pf*β-tubulin was kept at 10 μM.

### 3.5 Virtual screening of MMV676477 and its analogs against *Pf*αI/*Pf*β-tubulin heterodimer

To test the hypothesis that the parent scaffold, MMV676477 and its analogs (n = 29) confer their biological activities by their interaction with either of the *Pf*tubulin isoforms (αI and β), *fast docking* of the compounds against the generated homology modeling-based structural model of *Pf*αI/*Pf*β-tubulin heterodimer was done to computationally screen for ligands with propensity to bind pharmacologically relevant binding-pocket conformations (**Figure 1C**) of the protein. Compounds were arranged on the basis of their most negative free binding energies or binding affinity to the target binding-pocket, and subsequently ranked by decreasing value of their negative binding energies, *ΔG*_bind_ (kCal/mol), along the Y-abscissa (**Figure 2A**). Despite the high structural similarity among MMV676477 and its analogs, as evaluated by structural superimposition using BIOVIA Discovery Studio Visualizer v20.1.0.19295 (**Figure 1A**), significant differences in their *in silico* interaction behavior with *Pf*αI/*Pf*β-tubulin heterodimer were observed, which has been graphically represented in the form of an association curve (Pearson’s correlation coefficient, r = 0.5), with negative binding energies, *ΔG*_bind_ (kCal/mol), along the Y-abscissa and overlay similarity (with reference to the parent scaffold, MMV676477), along the X-abscissa (**Figure 2B).** The parent scaffold, MMV676477 was found to fit into the pharmacologically relevant overlapping binding pocket located at the interface of *Pf*αI/β monomers and interact via hydrophobic (predominantly) and polar contacts (H-bonds) with amino acid residues of the protein.

**Figure 2:**
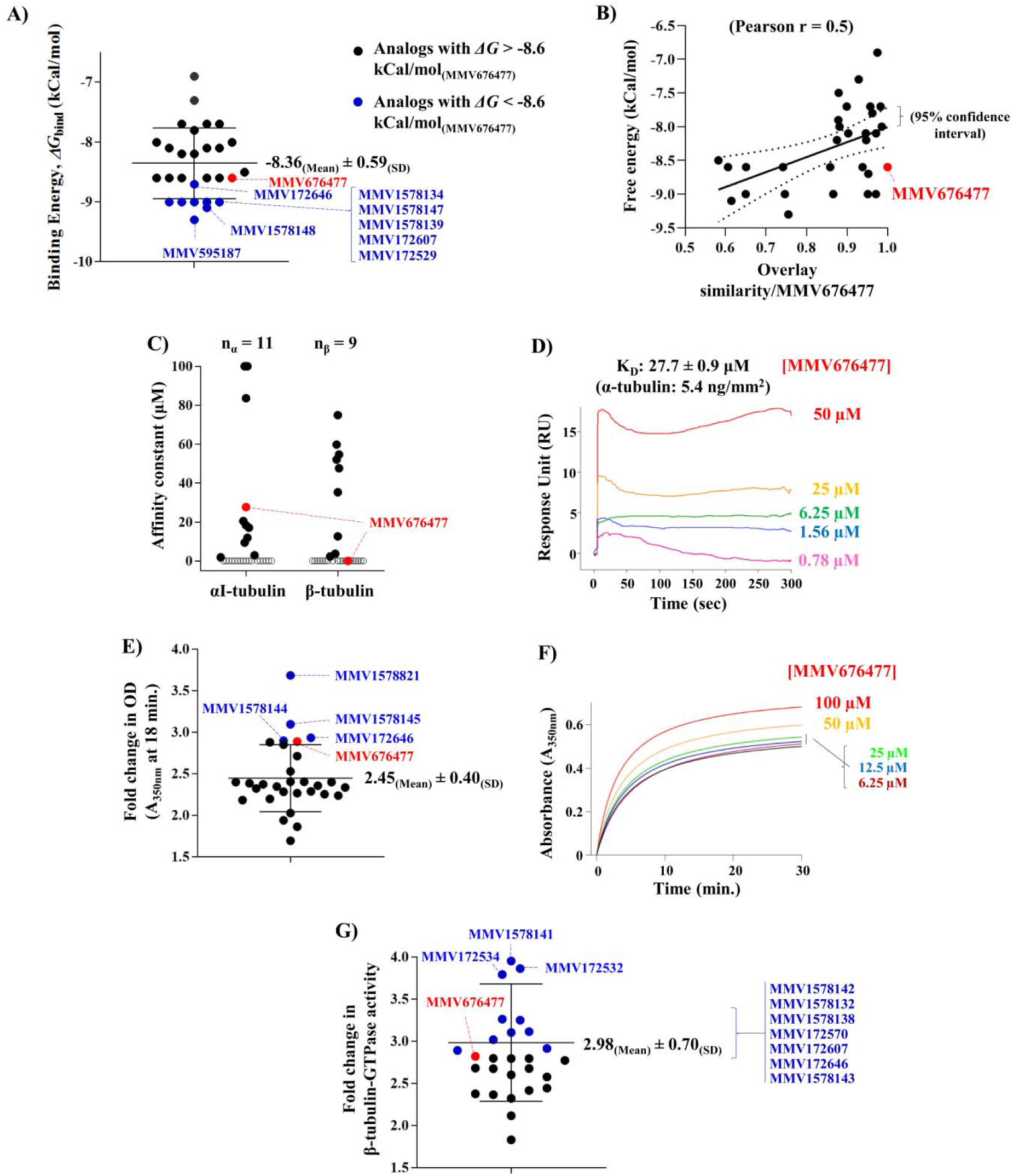
**Virtual screening of MMV676477 and its analogs against *Pf*αI/*Pf*β-tubulin heterodimer.** Compounds were arranged on the basis of their most negative free binding energies or binding affinity to the target binding-pocket, and subsequently ranked by decreasing value of their negative binding energies, *ΔG*_bind_ (kCal/mol), along the Y-abscissa **(a)**. Despite the high structural similarity among MMV676477 and its analogs, significant differences in their *in silico* interaction behavior with *Pf*αI/*Pf*β-tubulin heterodimer were observed, which has been graphically represented in the form of an association curve, with negative binding energies, *ΔG*_bind_ (kCal/mol), along the Y-abscissa and overlay similarity (with MMV676477), along the X-abscissa **(b). MMV676477 and its analogs interact with *Pf*tubulins and disturb *Pf*microtubule assembly dynamics.** SPR-based interaction analysis indicated that MMV676477 and its analogs displayed a wide range of binding affinities for *Pf*αI and/or *Pf*β-tubulin. Compounds were arranged on the basis of their binding affinities to the target binding-pocket, and subsequently ranked by increasing value of their affinity constants, K_D_ (μM), along the Y-abscissa **(c)**. A representative concentration dependent real-time sensorgram for the interaction of the parent scaffold, MMV676477 with *Pf*αI-tubulin is also shown **(d). Inhibitory effect (if any) on *Pf*microtubule assembly and GTPase activity conferred by *Pf*β-tubulin, in a cell free system.** Purified *Pf*αI- and *Pf*β-tubulins were incubated with either MMV676477 or its analogs and the absorbance at 350 nm was recorded as an indicator of tubulin polymerization. Compounds were arranged on the basis of their fold change in OD_350nm_, with respect to MMV676477, and subsequently ranked along the Y-abscissa **(e)**. MMV676477 and its analogs promoted *Pf*microtubule polymerization in a time-dependent manner, particularly MMV1578821, MMV1578144, MMV1578145 and MMV172646, showing fold-change in OD_350nm_ relatively more than MMV676477. A representative curve for the parent scaffold, MMV676477 is shown in **(f)**. We further investigated GTPase activity in the reaction mixtures prepared for the *in vitro Pf*microtubule assembly assay, at the end time-point, *i.e.*, 18 minutes. Compounds were arranged on the basis of their fold change in β-tubulin-GTPase activity, with respect to MMV676477, and subsequently ranked along the Y-abscissa **(g)**. The results showed that MMV676477 and its analogs promoted β-tubulin-GTPase activity, particularly MMV1578141, MMV172532, MMV172534, MMV1578142, MMV1578132, MMV1578138, MMV172570, MMV172607, MMV172646 and MMV1578143, showing fold-change in β-tubulin-GTPase activity relatively more than MMV676477.

### 3.6 MMV676477 and its analogs interact with *Pf*tubulins and disturb *Pf*microtubule assembly dynamics

To validate a possible mechanism of action of MMV676477 and its analogs *in vitro*, and to further validate the *in silico* interaction analysis with recombinant *Pf*αI and *Pf*β-tubulins (6X-His at their N’-termini), SPR-based interaction analysis was performed for the binding analysis, as mentioned in the methods section. Once immobilized on the SPR sensor chip surface, *Pf*αI and *Pf*β-tubulins demonstrated good stability throughout the experiment. For the potent binders, with an increase in mass concentration of the compounds, a gradual increase in SPR sensor signal was observed which linearly correlated with a corresponding change in refractive index of the medium immediately adjacent to the SPR sensing surface. MMV676477 and its analogs displayed a wide range of binding affinities for *Pf*αI and/or *Pf*β-tubulin. Compounds were arranged on the basis of their binding affinities to the target binding-pocket, and subsequently ranked by increasing value of their affinity constants, K_D_ (μM), along the Y-abscissa (**Figure 2C**). A representative concentration dependent real-time sensorgram for the interaction of the parent scaffold, MMV676477 with *Pf*αI-tubulin is shown in **Figure 2D**. Notably, MMV676477 showed significant interaction with *Pf*αI-Tubulin (K_D_ value of 27.7 ± 0.9 μM), but not with *Pf*β-tubulin.

SPR analysis suggests that most of the compounds are capable of interacting with either *Pf*αI or *Pf*β-tubulin, or both, indicating their probable tendency to disturb *Pf*microtubule assembly dynamics that could be an early event leading to mitotic arrest and/or morphological changes in the parasite. To verify this inference *in vitro*, the compounds were further examined for their inhibitory effect (if any) on *Pf*microtubule assembly and GTPase activity conferred by *Pf*β-tubulin, in a cell free system.

Purified *Pf*αI- and *Pf*β-tubulins were incubated with either MMV676477 or its analogs and the absorbance at 350 nm was recorded as an indicator of tubulin polymerization. Compounds were arranged on the basis of their fold change in OD_350nm_, with respect to the parent scaffold, MMV676477, and subsequently ranked along the Y-abscissa (**Figure 2E**). The results showed that MMV676477 and its analogs promoted *Pf*microtubule polymerization in a time-dependent manner, particularly MMV1578821, MMV1578144, MMV1578145 and MMV172646, showing fold-change in OD_350nm_ relatively more than MMV676477. A representative curve for the parent scaffold, MMV676477 is shown in **Figure 2F**. As an assay-positive control, paclitaxel was used which enhanced polymerization (Data not shown). These results demonstrate that MMV676477 and its analogues can interact with *Pf* tubulin(s) directly and inhibit *Pf*microtubule polymerization *in vitro*.

β-tubulin possesses GTPase activity, and the GTP hydrolysis changes the confirmation of tubulin molecules, thus driving the dynamic behavior of microtubules. Therefore, to test directly whether the parent scaffold, MMV676477 and its analogs inhibit or enhance β-tubulin-GTPase activity, we investigated GTPase activity in the reaction mixtures prepared for the *in vitro Pf*microtubule assembly assay, at the end time-point, *i.e.*, 18 minutes, using Malachite Green based assay, as described in the methods section. Compounds were arranged on the basis of their fold change in β-tubulin-GTPase activity, with respect to the parent scaffold, MMV676477, and subsequently ranked along the Y-abscissa (**Figure 2G**). The results showed that MMV676477 and its analogs promoted β-tubulin-GTPase activity, particularly MMV1578141, MMV172532, MMV172534, MMV1578142, MMV1578132, MMV1578138, MMV172570, MMV172607, MMV172646 and MMV1578143, showing fold-changes in β-tubulin-GTPase activity relatively more than MMV676477. As an assay-positive control, paclitaxel was used which enhanced β-tubulin-GTPase activity (Data not shown). These results demonstrate that MMV676477 and its analogs, indeed, interact with *Pf*tubulin(s) directly and disturb *Pf*microtubule assembly (polymerization and β-tubulin-GTPase activity) *in vitro*.

### 3.7 Screening the MMV tubulin targeting compounds for antimalarial activity against human malaria parasite

Microtubules are integral parts of eukaryotic cells. In malaria parasites they serve a variety of functions at different stages of their life cycle. During asexual erythrocytic stage of its life cycle, microtubules are very much required during nuclear division, partition of the organelle and cytosol into new merozoite and invasion of erythrocyte by merozoite [4,59–61].

MMV tubulin targeting compounds from the pyrazolopyrimidine series were screened for their antimalarial activity against asexual stage of human malaria parasite *in vitro*. It is imperative that new antimalarial drugs undergo cross-resistance studies, to ensure that they can withstand current parasite resistance mechanisms in order to ensure their clinical effectiveness. All compounds were screened for their antimalarial activity against RKL-9 and R539T strains of *P. falciparum*.

For an IC_50_ determination, highly synchronized ring-stage parasites of 3D7, RKL-9 and R539T were exposed to a range of compound concentration respectively for 72 h. Assays were performed in 96-well plates, with a maximum in-well DMSO concentration of 0.35%. Parasitemia was measured at 72 h post-treatment using SYBR Green I based fluorescence assay and graph was plotted for the average value of three independent sets of experiments **(Figure 3a (i-iii))**. Fifty percent parasite growth inhibition was reserved as a cut-off value for the selection of compounds to have anti-plasmodial activity. Data revealed a potent anti-malarial activity of MMV compounds against 3D7, RKL-9 and R539T strain of *P. falciparum* with a significant reduction in the parasite load, with IC50 value ranging from ∼150 nM to 5 µM. Among the 30 compounds tested, three compounds MMV676477, MMV1578136, MMV1578138 showed higher potency across all the strain tested (3D7, RKL-9 and R539T) with IC_50_s of 0.54 µM, 0.95 µM, 0.66 µM in 3D7, 0.625 µM, 0.725 µM, 0.520 µM in RKL9 and 0.800 µM, 0.25 µM and 0.27 µM in R539T respectively. The IC_50_ values of the *P. falciparum* 3D7 drug sensitive line were compared with the IC_50_ against drug resistant lines (RKL-9 and R539T) to calculate a Resistance Index (RI; resistant line IC_50_/sensitive line (3D7) IC_50_; the higher the RI, the higher the resistance level). RI’s were compared for assays carried out over 72 h, antimalarial drugs chloroquine, and artemisinin were used as positive controls for RKL-9 and R539T, respectively (need to be calculated). Compounds MMV676477, MMV1578136 and MMV1578138 RI showed no cross-resistance against RKL-9 with an RI of 1.157, 0.76, 0.7 respectively. Similarly, RI against R539T was coming out to be 1.48, 0.26 and 0.409 respectively. Observed low levels of cross-resistance indicated that pre-existing resistance may not be an impediment to the development of these drugs as antimalarials.

**Figure 3:**
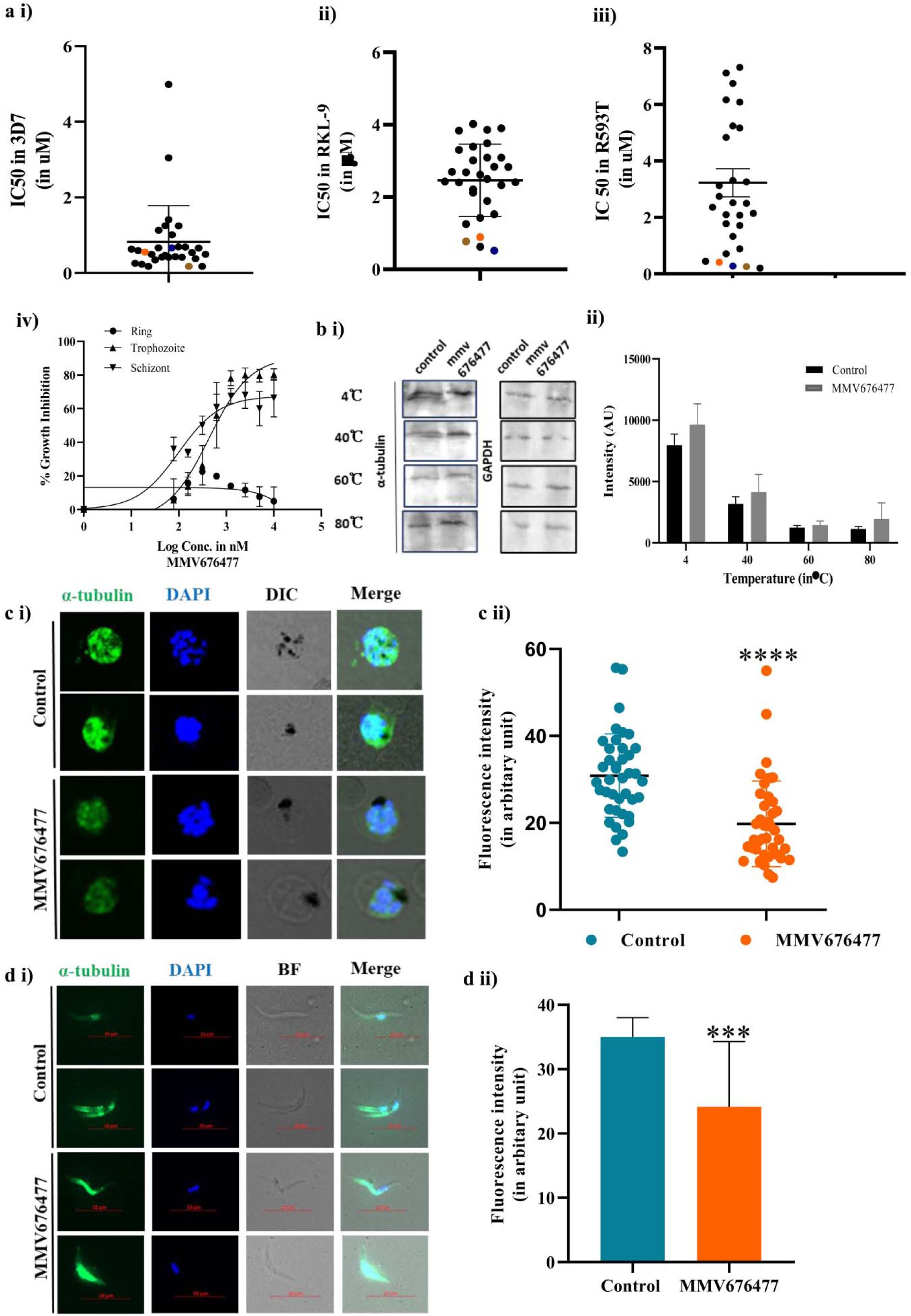
**(a) *In vitro* growth inhibition assay in *P. falciparum* with MMV tubulin targeting compound. (i)** Graph represent the IC50 in 3D7 (Chloroquine sensitive line) strain of malaria parasite with all the MMV compounds tested **(ii)** represent the IC50 obtained in RKL-9 (Chloroquine resistant line) strain of malaria parasite **(iii)** represent the IC50 obtained in R539T (Artemisinin resistant line) strain of malaria parasites (MMV676477 in (), MMV1578136 in () and MMV1578138 in () **(iv)** Graph represent dose response curve obtained with compound MMV676477 when treated at ring (0-6 h), trophozoite (22-26 h) and schizont stage (38-42 h) of P. falciparum (3D7) respectively for 10 h . IC50 was calculated using non-linear regression curve analysis in GraphPad prism 8.0 software. Data represent the mean ± S.D of three biological replicate. **(b) CETSA based target identification of MMV676477 in malaria parasite cell lysate. (i)** Figure depicts the immune-based CETSA investigation of Pf alpha-tubulin I thermostability in the presence and absence of MMV676477. Saponized parasite lysates were lysed and parasite protein was pre-incubated for 30 minutes with MMV676477 (10M), whereas parasite protein was treated with DMSO as a control. After thermal denaturation at 40, 60, and 80°C, samples were examined by western blot using an alpha-tubulin antibody. The intensity of the protein at 4°C was used as a control. **(ii)** Densitometry quantification of the respective immunoblot was performed using ImageJ software and plotted as bar graph.. Blotting with anti-GAPDH served as loading control. Data represent the mean ± S.D of n=2. **(c) c) Fluorescence light microscopy image of microtubule during blood stage schizogony. (i)** *P.falciparum* at schizont stage were treated with MMV676477 for 6 h at IC50 concentration. Following treatment, cells were pelleted down, and smear was made that was subsequently stained for alpha-tubulin (green) and image was captured under 100X oil objective on a Nikon Ti2 microscope. Fluorescence microscopic images represent the alpha-tubulin staining in DMSO and compound treated parasite respectively. Nuclear body stained with DAPI and DIC were shown. **(ii)** Graph shows comparative intensity profile of alpha-tubulin in 40 different RBCs with and without MMV 676477 treatment. Error bar represent mean ± S.D with P-value < 0.0001 ****. **(d) MMV compound Binds at the alpha-beta tubulin dimer interface and promote tubulin polymerization at sporozoite stage (i)** Sporozoite treated for 1 h with 3 µM of MMV676477 were fixed in chilled methanol and stained for alpha-tubulin and analyzed under fluorescence microscope (Carl Zeiss Microscopy, LLC at 100X objective). Fluorescence microscopic image showed enhanced microtubule assembly in treated sporozoite when compared to DMSO treated sporozoite. **(ii)** Graph on the right shows comparative intensity profile of alpha-tubulin in 40 different sporozoite with and without MMV compound treatment. Error bar represent mean ± S.D P-value < 0.001 ***

### 3.8 Asexual Blood Stage specificity profile of MMV676477 against human malaria parasite

A previous study using a variety of microtubule inhibitors has shown that the susceptibility of the parasite to these inhibitors varies with the developmental stage of the parasite [62, 63]. For instance, dolastatin 10, vinblastine and Taxol demonstrated great efficacy when applied to cultured schizont stages, less effective when applied to trophozoites, and were ineffective when applied to rings [64].

To examine the susceptibility of these compound on distinct stages of *P. falciparum* intra-erythrocytic development [6], *in vitro* stage and concentration-dependent effects of MMV676477, MMV1578136 and MMV1578138 were studied with synchronous cultures of *P. falciparum* 3D7. In this assay synchronized ring (0-6 h), trophozoite (22-26 h) and schizont stages (38-42 h) were exposed to different concentration of drug (20, 10, 5, 2.5, 1.25, 0.625, 0.312, 0.156 µM) for 10 h. Post 10 h of exposure, compound was removed by extensive washing and allowed to grow into the next cycle. Parasites growth was determined using SYBR green I based fluorescence assay. Half-maximal inhibitory concentrations (IC_50_) were derived by non-linear regression analyses of the dose-response data. The IC_50_ values based on these 10 h exposures at specific asexual blood stages is referred as IC_50_10 h. Result showed that IC_50_10 h for MMV676477 at ring, trophozoite and schizont stages were > 20 µM, 0.431 µM and 0.098 µM respectively **(Figure 3a (iv))**. Similarly for MMV1578136 and MMV1578138, the IC_50_10h for ring, trophozoite and schizont stage is coming out to be >10 uM, 457 nM, 335 nM, and >10 uM, 299 nM, 159.4 nM respectively. This suggests stage specific inhibitory effects of these compound on *P. falciparum* growth. In contrast to 72 hours of exposure, the parasite treated during the schizont stage had a lower IC50, which is presumably due to the presence of a larger population of drug-susceptible parasites. These results may be explained by the fact that tubulin expression is lowest in the ring, maximum in the trophozoite, and highest in the schizont. However, there is also a plausible alternative explanation, which suggests that ring-stage parasites are simply less permeable to the inhibitors. A set of intranuclear spindle microtubules orchestrates nuclear division during schizogony [15, 16]. Consequently, these compounds appear to be primarily acting on the mitotic microtubules and inhibiting nuclear division by stabilizing the microtubular structures associated with the mitotic apparatus. Also, during erythrocyte invasion, the torsion/contraction of subpellicular microtubules is very much needed for merozoite motility [65]; disturbing dynamicity of microtubules could also impact this process.

### 3.9 Confirmation of MMV676477 binding with Plasmodium tubulin by Cellular Thermal Shift Assay (CETSA)

It is imperative to identify targets in order to understand the mechanisms of action of new therapeutics. To validate or confirm that MMV676477 binds to the *P. falciparum* tubulin, conventional immuno-based CETSA was performed [52, 53]. A lysed saponized parasite pellet was incubated with 10 µM of MMV676477 for 30 min at RT. The protein fractions were then subjected to a panel of temperature ranging from 4-80 ℃ for 3 min, followed by cooling down in ice for 3 min so that the potential thermal stabilization could be assessed. Each of the fractions were then centrifuged at 10,000 rpm for 30 min. Supernatant fractions were then loaded on SDS-PAGE gel and immunoblot analysis were performed with anti-alpha tubulin antibody to confirm the stability of this protein in the presence of its inhibitor. Western blot analysis revealed the stability of protein in the presence of its binding molecule. At 40℃ there is no difference in the baseline of protein amount in the presence or absence of inhibitor, while increasing the temperature gradually decrease the stability of protein in absence of MMV676477, indicating that thermal stability of alpha-tubulin is affected by the presence of its targeting molecule **(Figure 3b (i)).** Blotting with anti-GAPDH served as a loading control. Simultaneously mean intensity of immunoblot at different temperatures was calculated by ImageJ software and plotted as bar graph as shown in **(Figure 3b (ii)).**

### 3.10 Effect of microtubule targeting compound on Plasmodium microtubule structure

Microtubules are dynamic cytoskeletal polymers that spontaneously switch between phases of growth and shrinkage. To study the effect of these microtubule targeting compound on microtubule dynamics of plasmodium parasite, as well as on sporozoite stage of plasmodium, Immunofluorescence analysis was performed. As expression of tubulin is higher at later asexual stage, there forth schizont stage parasite were treated with MMV676477 at IC50 concentration obtained from 72 h growth inhibition assay. Parasite were treated with compound for 6 h and processed further for immunofluorescence analysis. Similarly, to study microtubule dynamicity at sporozoite stage isolated sporozoite were treated with MMV676477 and processed further for immunofluorescence analysis using alpha-tubulin antibody. **Figures 1 c(i)** & **d(i)** shows immunofluorescence images of untreated and MMV676477 treated parasite as well as sporozoites. **Figure 1 c(ii)** and **Figure 1 d(ii)** represent the mean intensity plot of alpha-tubulin in schizont stage parasites as well as in sporozoites calculated as described in material and method sections

In untreated parasites, **(Figure 1 c(i))** microtubule structure formed by them was observed in association with the nucleus. When treated with MMV676477 diffused microtubule structure was observed. However contrasting result is observed in case of sporozoite where stabilized microtubule structure was observed when it was treated with MMV676477. Difference in observed result between asexual parasite and sporozoite could be due to the fact that treatment of schizont stage parasite with non-saturating concentration of microtubule stabilizing agent (∼500 nM) can result in microtubule catastrophe events in cell [66–69], which is also being observed in our immunofluorescence data. However, in case where saturable amount of MMV676477 is present it stabilizes the microtubule by cooperatively binding at the PfαI-β dimer interface thereby preventing microtubule shrinkage.

### 3.11 Cytotoxicity Analysis of microtubule targeting compounds in mammalian hepatocyte cell line (HepG2)

In order to determine for any potential real-life application of our compounds, the cytotoxicity of all the thirty compounds was investigated using mammalian hepatocyte cell line HepG2. The cytotoxicity (EC50 values) of these complexes were > 30 μM HepG2 cell lines indicating high cytotoxicity in parasitic cell lines in comparison to mammalian cells. Selective indices (SI) were also calculated for all the compounds and the result showed higher selectivity towards malaria parasite over mammalian cells, suggesting they can be used as a potential antimalarial agent (**Table 1**). Out of all the thirty compounds tested MMV676477, MMV1578136 and MMV1578138 showed higher selective index as well as the potent antimalarial activity with low RI. As a result, they are chosen for further investigation.

**Table 1:** Summary of IC50, Resistance Indexes (RI) and Selectivity Indexes (SI) of all thirty compounds.

| **MMVCompound** | **3D7** | **RKL9** | **R593T** | **RI(RKL9-3D7)** | **RI(R593T-3D7)** | **HepG2** | **SI** |
| --- | --- | --- | --- | --- | --- | --- | --- |
| **MMV001212** | 0.69 | 2.50 | 2.50 | 3.63 | 3.64 | 14.11 | 20.50 |
| **MMV001213** | 1.13 | 0.63 | 2.08 | 0.55 | 1.84 | 15.59 | 13.78 |
| **MMV1578131** | 0.54 | 1.25 | 2.52 | 2.30 | 4.64 | 7.083 | 13.04 |
| **MMV1578132** | 0.63 | 3.30 | 1.33 | 5.23 | 2.10 | 27.4 | 43.49 |
| **MMV1578133** | 0.42 | 2.43 | 7.31 | 5.83 | 17.56 | 8.414 | 20.21 |
| **MMV1578134** | 0.18 | 2.12 | 0.44 | 11.84 | 2.48 | 24.4 | 135.55 |
| **MMV1578135** | 1.41 | 1.42 | 10.83 | 1.01 | 7.67 | 15.56 | 11.01 |
| **MMV1578136** | 0.18 | 0.78 | 0.26 | 4.33 | 1.43 | 71.88 | 401.56 |
| **MMV1578137** | 1.25 | 3.49 | 2.14 | 2.80 | 1.72 | 36.71 | 29.43 |
| **MMV1578138** | 0.67 | 0.52 | 0.28 | 0.78 | 0.41 | 60.92 | 91.19 |
| **MMV1578139** | 0.23 | 2.62 | 3.12 | 11.28 | 13.47 | 42.01 | 181.07 |
| **MMV1578140** | 0.42 | 3.84 | 2.75 | 9.11 | 6.52 | 29.7 | 70.46 |
| **MMV1578141** | 0.50 | 1.89 | 4.83 | 3.80 | 9.70 | 12.41 | 24.91 |
| **MMV1578142** | 0.59 | 3.90 | 2.36 | 6.59 | 3.98 | 6.26 | 10.58 |
| **MMV1578143** | 0.47 | 3.09 | 1.78 | 6.58 | 3.78 | 3.86 | 8.22 |
| **MMV1578144** | 0.69 | 3.01 | 3.26 | 4.34 | 4.70 | 23.5 | 33.86 |
| **MMV1578145** | 1.01 | 2.83 | 2.10 | 2.79 | 2.07 | 7.48 | 7.38 |
| **MMV1578146** | 0.26 | 3.86 | 5.16 | 15.09 | 20.20 | 41.04 | 160.50 |
| **MMV1578147** | 0.42 | 2.68 | 6.17 | 6.46 | 14.84 | 2.22 | 5.35 |
| **MMV1578148** | 0.66 | 2.41 | 3.14 | 3.63 | 4.74 | 18.05 | 27.22 |
| **MMV1578821** | 0.35 | 2.40 | 6.09 | 6.89 | 17.44 | 12.65 | 36.25 |
| **MMV172529** | 0.38 | 3.39 | 1.72 | 8.89 | 4.50 | 29.04 | 76.12 |
| **MMV172532** | 0.43 | 2.33 | 0.20 | 5.36 | 0.46 | 9.108 | 20.95 |
| **MMV172534** | 4.99 | 3.09 | 3.31 | 0.62 | 0.66 | 28.28 | 5.66 |
| **MMV172570** | 0.18 | 1.52 | 7.11 | 8.64 | 40.38 | 35.28 | 200.34 |
| **MMV172573** | 3.05 | 2.84 | 0.89 | 0.93 | 0.29 | 2.48 | 0.81 |
| **MMV172607** | 0.50 | 2.21 | 6.75 | 4.44 | 13.55 | 86.81 | 174.28 |
| **MMV172646** | 0.67 | 2.69 | 5.24 | 4.03 | 7.84 | 42.6 | 63.71 |
| **MMV595187** | 1.25 | 4.02 | 0.71 | 3.21 | 0.57 | 23.87 | 19.08 |
| **MMV676477** | 0.56 | 0.89 | 0.41 | 1.60 | 0.74 | 79.06 | 142.16 |

### 3.12 MMV676477, MMV1578136 and MMV1578138 inhibited the *P. berghei* sporozoites infection in human hepatocyte in vitro

In order to determine whether these compounds affect the pre-erythrocytic phase of the malaria parasite, sporozoite infection in human hepatoma cell line HepG2 was performed in the presence or absence of microtubule targeting compound MMV676477, MMV1578136 and MMV1578138 at 3 µM concentration. After 48 h, cells were collected followed by RNA isolation and cDNA synthesis. Real-time quantitative PCR analysis with *P. berghei* 18S primer was then performed to read out the sporozoite infectivity in the presence and absence of compound. The *in vitro* sporozoite infection assay showed 43, 72 and 65.5% decrease in 18S rRNA expression for compound MMV676477, MMV1578136 and MMV1578138 respectively when compared to control **(Figure 4a)**.

**Figure 4:**
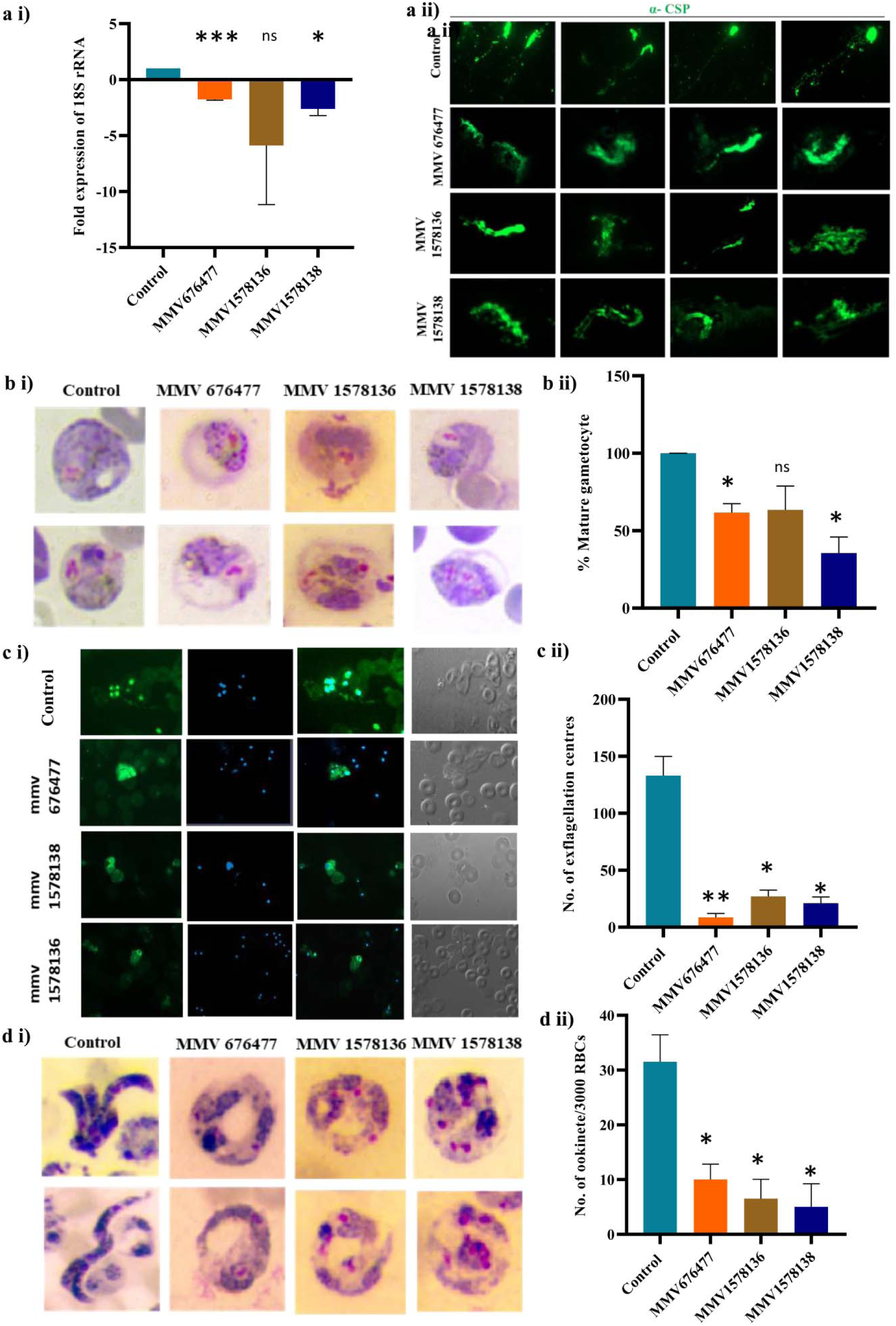
**(a i) MMV compound inhibits HePG2 infection by *Plasmodium berghei* sporozoites *in vitro***. Sporozoites isolated from salivary gland of anopheles mosquito were allowed to infect HePG2 (2× 10^5^) *in vitro* in presence of 3 µM of MMV compound (MMV676477, MMV1578136 and MMV1578138). After 48 h total RNA was extracted, cDNA was synthesized and *P.berghei* 18S rRNA was quantified by real-time PCR. Bar graph represent the percent decrease in expression of 18S rRNA in and treated culture incomparison to control. Mean value of relative 18S expression was normalized by human beta-actin expression. The graph was generated from three independent experiments done in triplicate. The error bars represent standard deviations for three experiments with P-value < 0.01**, P-value < 0.05* **(a ii) MMV compound showed potent inhibition of sporozoite motility.** Sporozoite motility assay was performed in presence of MMV676477, MMV1578136 and MMV1578138 at 3 µM concentration. Representative immunofluorescence images of WT and compound treated sporozoites, revealed with the anti-CSP antibody. DMSO treated sporozoites display the typical continuous gliding pattern, in contrast, compound-treated sporozoites are either non-motile or have discontinuous gliding patterns (green). **(b) Tubulin targeting MMV compound inhibit maturation of gametocyte. (i)** represent the micrograph of giemsa stained mature gametocyte in presence and absence of MMV compound. **(ii)** Cells were scored as either mature or immature gametocyte and plotted as a percentage mature gametocyte. The results are derived from a representative experiment performed in triplicate. Error bars represent mean ± S.D with P-value < 0.05 * **(c) Tubulin targeting MMV compound inhibit ex-flagellation of male gametocyte. (i)** Bar graphs represent the no of ex-flagellation centre observed in 10 field views following addition of MMV compound. Three MMV compounds (MMV676477, MMV1578136 and MMV1578138) with activity against *P.berghei* gametocyte were evaluated for their ability to inhibit male gamete ex-flagellation. Tail blood from *P. berghei* infected mice was mixed pre-warm complete RPMI. and pre-incubated with 3 µM of MMV676477, MMV1578136 and MMV1578138 separately for 1 h at 37°C. Ex-flagellation was induced by transferring the sample to 19°C in presence of 100 µM of xanthurenic for 10-12 min. sample were then smeared onto glass slide and processed further for immunofluorescence analysis using alpha-tubulin antibody. Images were captured at 100X objective using Carl Zeiss Microscopy, LLC. Fluorescence images represent the flagella (in green) coming out from the activated male gametocyte while no such exflagellated gametocyte were observed in treated samples. **(ii)** Bar graph represent the no of ex-flagellation center observed in 10 field views in each of the sample. For these induced gametocytes after 15 min of induction were aliquoted onto glass bottom plate and observed under bright field microscopy at 40X magnification Data represent the mean ± S.D (n=3) B) Mature gametocyte of treated and untreated culture after induction were stained for alpha-tubulin (green). Nuclei stained with DAPI and DIC were shown. **(d) MMV compound inhibit *ex vivo P*. *berghei* Ookinete development. (i)** Ookinete conversion assays were performed using *P. berghei* ANKA parasites. Blood collected from an infected mice were dispensed in 96 well plate containing ookinete media along with MMV676477, MMV1578136 and MMV1578138 for 24 h at 21°C. For control, parasites were treated with DMSO alone. After incubation, thin blood smear was prepared from each of the treated well, stained with giemsa and observed under light microscope. Fully mature, banana-shaped ookinetes observed in DMSO treated culture, Parasite treated with tubulin targeting compound failed to form mature ookinete **(ii)** Bar graph represent the percent of mature ookinete in DMSO control and treated culture. Data represent mean ± S.D. (n=3) with p-value < 0.05 **

The observed reduction in sporozoite infection could be due to the inhibition of several steps that comprise the infection process - sporozoite motility, host cell invasion, or intravacuolar development of the parasite [70]. As these compounds were believed to target microtubules, which are necessary for sporozoite motility and thus infectivity, sporozoite motility assays were conducted. To this end, sporozoites were isolated from freshly dissected mosquito salivary glands and allowed to glide on the slide surface in the presence or absence of MMV676477, MMV1578136 and MMV1578138 at 3 µM for 30 min. The Circumsporozoite Protein (CSP) trails shed by the sporozoites while gliding was probed with an anti- CSP antibody and observed under a fluorescent microscope [70–72]. Immunofluorescence analysis demonstrated the circular and zig-zag motion of sporozoite in DMSO treated control while no such motility was being observed in drug treated sporozoites **(Figure 4b)**. The result suggested inhibition in sporozoite motility could be the possible reason behind decreased parasite infection in hepatocyte.

### 3.13 MMV676477, MMV1578136 and MMV1578138 inhibited the gametocyte rounding-up of *P. berghei* parasite *ex-vivo*

To effectively eradicate malaria, a drug must be effective against both the asexual blood stages as well as the sexual stages. Gametogenesis involves rounding up of activated gametocytes, ex-flagellation of male gametes, and gamete egress. Microtubules are disassembled both in gametocyte maturation and during cell division. In the final phase of gametocyte maturation, microtubules depolymerize, causing the parasite to shorten and the red blood cells to become more deformable [73–75]. Because microtubule disassembly is required for the maturation of gametocyte and MMV676477 previously has been shown to stabilize microtubule assembly [76]. We next hypothesized that these compounds could be targeting the rounding up of gametocyte.

To evaluate the gametocyte, round up and egress from the RBC blood was obtained from *P. berghei* infected mice and incubated with test compounds for 1 h at 3 µM concentration. Gametocyte induction was given in presence of ex-flagellation media containing 100 µM xanthurenic acid at 20°C. Within 15 minute of gametocyte induction Giemsa smears were prepared and observed under light microscope **(Figure 4c (i))**. In MMV676477, MMV1578136 and MMV1578138 treated culture the numbers of mature rounded up gametocyte were reduced by approximately half indicating gametocyte maturation was affected. While in DMSO treated parasite mature round shaped gametocyte was observed as shown in **figure 4c (ii)**.

### 3.14 MMV676477, MMV1578136 and MMV1578138 inhibit ex-flagellation of *P. berghei* parasites *ex vivo*

Microtubule dynamics are crucial for the ex-flagellation of male gametocytes. Axonemes microtubules are responsible for the flagellar movement in male gametes. During ex-flagellation, male gametes produce eight highly motile, slender microgametes, which penetrates the macrogamete and produce a zygote [21, 22]. Motile machinery of the sexually active gametes is comprised of flagella that are primarily composed of microtubules, since tubulin is the major protein component of microtubules, it is likely that these tubulin-targeting compounds could affect the highly motile process of ex-flagellation. Compounds that disrupt the formation of male and/or female gametes will further interfere with the downstream process which in turn disrupt malaria transmission.

To assess the effect of MMV676477, MMV1578136 and MMV1578138 on male ex-flagellation, we carried out *ex vivo* exflagellation assay with *P. berghei* infected RBCs obtained from infected mice. For this *P.berghei* infected mice blood was pre-incubated with the mentioned compound for 1 h at concentration of 3 µM at 37°C. After incubation with the tested compound, ex-flagellation was induced by the addition of 100 µM xanthurenic acid at RT for 15 min. Afterwards, the number of ex-flagellation centers was microscopically counted in both DMSO and compound treated parasites at 40X magnification. Ex-flagellation is a highly time-dependent characterized by the emergence of the male gametes from the infected erythrocyte after a period of approximately 15 to 20 minutes. Once they emerge from the gametocyte residual body, they migrate away from the ex-flagellation center through flagellar locomotion and continue to move until either they reach a female gamete and fertilize it. It is easy to distinguish male gametes by light microscopy since they are highly motile cells. In a monolayer of erythrocytes, ex-flagellation cause local disturbance of nearby cells while leaving distant cells motionless. All the tested compounds showed a significant reduction in the numbers of ex-flagellation centers by approximately 80% in comparison to DMSO control **(Figure 4d (ii))**.

To further assess the defect in ex-flagellation upon treatment, IFA with α-tubulin antibody was performed with the DMSO treated and compound treated parasite after induction of gametogenesis. Labeling of the male gamete with α-tubulin antibody showed flagellar motile structures comprised of axoneme microtubule while no such flagellated male gamete was observed in parasite treated with compound MMV676477, MMV1578136 and MMV1578138 **(Figure 4d (i))** suggesting targeting microtubule main component of the cytoskeleton, leads to the inhibition of ex-flagellation by interfere with the formation of flagella.

### 3.15 Effect of MMV676477, MMV1578136 and MMV1578138 on ookinete maturation of *P. berghei* parasite *ex vivo*

In malaria-causing parasites, a round zygote develops into an elongated motile ookinete within the stomach of a mosquito. Round zygotes undergo major and rapid morphological changes. Microtubules play an important role during the initial stages of zygote to ookinete formation. During ookinete maturation zygote grow in length, and mature into elongated, crescent-shaped ookinetes [24]. The cytoskeletal structures of the pellicle are responsible for this process, particularly the inner membrane complex (IMC) and subpellicular microtubules (SPMs). These subpellicular microtubule network consists of longitudinal arrays of alpha and beta-tubulin heterodimers that are crosslinked to one another and to the plasma membrane by various microtubule-associated proteins. The ookinete’s apical rings are formed at the same time and organize the SPM of > 40 microtubules [77, 78]. Taxol (microtubule polymerization agent) and colchicine (microtubule depolymerization agent) were found to inhibit the ookinete maturation. To evaluate the effect of MMV676477, MMV1578136 and MMV1578138 on gametocyte to ookinete maturation compounds were tested against *P. berghei* malaria parasite *ex vivo*. *P. berghei* infected RBCs were collected from infected mice and kept in ookinete culture media in presence of these compound at concentration of 3 µM for 24 h at 26 °C. After 24 h post-treatment, Giemsa smears were made and observed under light microscope **(Figure 4e (i))**. Ookinete numbers were counted in each of the untreated and treated groups, and represented as a bar graph **(Figure 4e (ii))**. Fully developed elongated morphology of ookinete was observed in the DMSO treated control group, while morphologically immature or partially developed ookinetes were observed in the treated group.

Importantly, *in vitro* assessment of the stability of the parent scaffold, MMV676477 against mouse and human microsomes, LogD value, mouse and human liver hepatocytes, effectibve permeability and sustained plasma stability prove that the compound possesses excellent overall stability (**Table 2**).

**Table 2:**
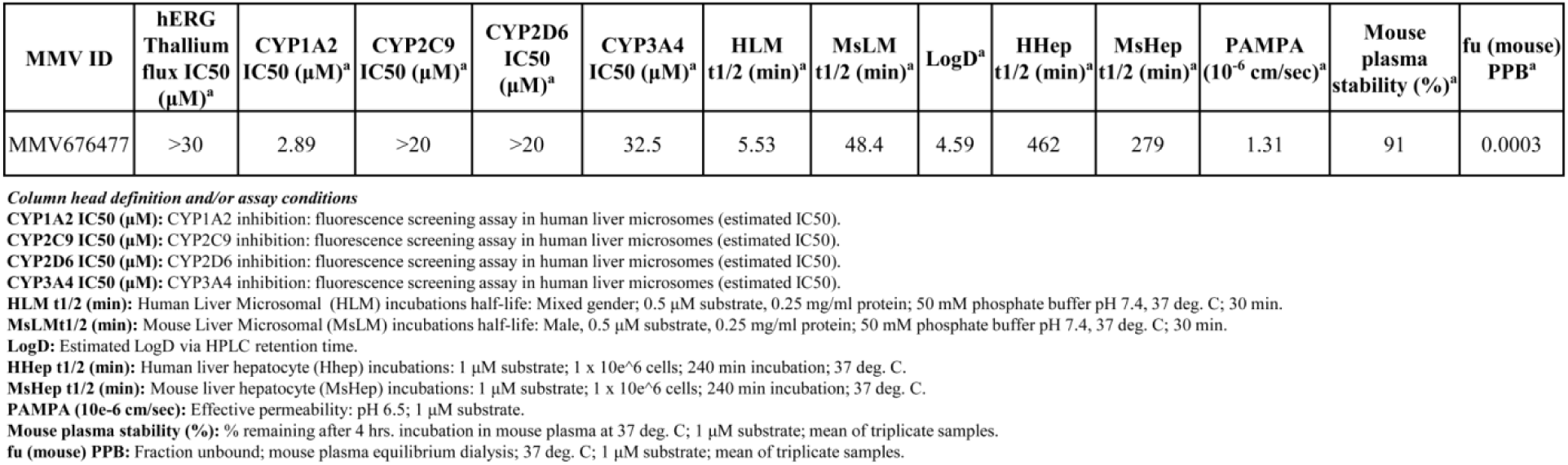
Summary of *in vitro* ADME/T properties of the parent scaffold, MMV676477.

## 4. DISCUSSION

The next generation of antimalarials will have a preference for molecules that have multiple stages of activity. However, these molecules are not always discovered in diversity library screens. Several effective antimalarial compounds have been discovered over the past decade which may serve as starting points for new treatments against malaria. To counter growing resistance to traditional therapy regimens, small molecules with distinct mechanisms of action for killing *Plasmodium* parasites are being investigated.

Microtubules, which have recently received considerable attention for the development of anticancer therapies, could also be efficient antimalarial targets. Cell division, mobility, and vesicular transport are all regulated by the microtubules of the cytoskeleton, whose key structural component is tubulin, which is potentially targeted by a wide variety of natural and synthetic drugs known as Microtubule-Targeting Agents (MTAs) [1]. These agents have been explored in the case of malaria as well. Vinblastine and Taxol, for instance, have been shown to be highly effective against malarial parasites that disrupt microtubule structures inside the erythrocyte, in relevant quantities [59,60,64]. Despite the fact that these inhibitors are useful, their promise as antimalarial medicines is limited by their significant toxicity to human cells. Furthermore, dinitroanilines and phosphorothioamidates, other microtubule inhibitors, exhibit low toxicity to mammalian cells, but have moderate activity against human malaria parasites [61,62,79]. Drugs that have higher selectivity for the malaria parasite and low toxicity towards mammalian cells will be preferred.

In this study, we provide a first report on the multistage potency of MMV676477 scaffold and its synthetic analogs, against intra-erythrocytic asexual, sexual and liver stage *P. falciparum* parasites *in vitro*. The activity profile of these compounds demonstrated their multistage-active antimalarial potency, which will prove to be extremely useful for eradicating malaria. Being from the same scaffold, the received compounds were first analyzed for their degree of structural similarity. *In silico* analysis using BIOVIA Discovery studio Visualizer tool, revealed that the majority have an overlay similarity index of more than 0.8. Compounds with similar structural architecture tend to have similar bioactivity profile. To test the same, all of these thirty compounds were screened against chloroquine resistant (RKL9, field isolate) and artemisinin resistant (R539T) cell lines as well. Among the thirty compounds screened, MMV676477, MMV1578136 and MMV1578138 showed potent anti-malarial activity with an IC_50_ of 0.54 µM, 0.95 µM, 0.66 µM in 3D7; 0.625 µM, 0.725 µM, 0.520 µM in RKL9; and, 0.800 µM, 0.25 µM and 0.27 µM in R539T line of *P. falciparum*, respectively, *in vitro*. Interestingly, no cross-resistance was identified with MMV676477, MMV1578136 and MMV1578138 among these parasite lines, implying their possible distinct mode of actions. Additionally, investigation of compound’s peak activities in stage-specific assay revealed that they are particularly active against late stages of asexual intra-erythrocytic development. As the parasite develops from the ring stage to trophozoite stage, and further to schizont stage, the expression of tubulin increases which may explain potent activity of these compounds at the later stages. Each individual nucleus in the schizont stage has its own Microtubule Organizing Center (MTOC), which is responsible for fulfilling its own microtubule nucleation requirements. Therefore, drugs that target microtubule dynamics at this point may have an adverse effect not only on the asexual development of the parasite, but also on the invasion of erythrocytes by newly formed merozoites. Cytotoxic effects of these derivatives further showed low cytotoxicity against mammalian liver cell line, HepG2. Due to its high toxicity towards *Plasmodium* and low cytotoxicity to mammalian cells, these compounds represent an exciting opportunity for the development of potent anti-malarials.

Recently, MMV676477 has been shown to inhibit Leishmania replication by stabilizing microtubule assembly [24]. To examine whether the same phenomenon occurs in *Plasmodium*, MMV676477 and its analogues were first screened virtually against the *Pf*αI/*Pf*β-tubulin heterodimer. Homology modelling-based structural model of *Pf*αI/*Pf*β -tubulin heterodimer was used to perform fast docking of drugs against the protein to screen for compounds that could potentially bind pharmacologically relevant binding-pocket conformations of the protein. Structural superimposition using BIOVIA Discovery Studio Visualizer v20.1.0.19295 showed that MMV676477 and its analogues high structural resemblance, although their *in silico* interactions with *Pf*αI/*Pf*β -tubulin heterodimer were found to differ dramatically (**Figure 2B**).

Further, *in vitro* analysis was performed to validate the *in silico* interaction analysis. SPR-based biochemical investigation indicated that the majority of the compounds are capable of interacting with *Pf*αI and/or *Pf*β-tubulin, indicating their probable potential of disrupting the dynamics of *Pf*microtubule assembly, which could be a precursor for mitotic arrest and/or morphological alterations in the parasite. To confirm this *in vitro*, the compounds were subsequently investigated in a cell free system for their inhibitory effect (if any) on *Pf*microtubule polymerization and GTPase activity imparted by *Pf*β-tubulin. *In vitro* polymerization experiments showed that these compounds are capable of enhancing the microtubules assembly (**Figure 2F**). Since, GTP molecules get rapidly hydrolyzed following incorporation into the polymerizing microtubule, GTPase activity of β-tubulin was performed via Malachite Green based assay. As shown in **figure 2G,** these compounds demonstrated they disturb the microtubule assembly dynamics by enhancing its polymerization.

Compounds found to enhance microtubule assembly *in vitro* were then screened for their tubulin polymerization property inside the cell. Towards this, immunofluorescence analysis with alpha-tubulin antibody was performed with MMV676477 treated schizonts as well as sporozoites. As observed under the fluorescence microscope (**Figure 3d(i)**), MMV676477 treatment of sporozoites stabilized microtubule structure, unlike the scattered microtubule structure that was observed in the case of schizont.. These findings were attributed to an understanding of how microtubule stabilizing agents behave when they are present in non-saturatable amounts. However, when they are present in saturatable amounts and accumulate at the binding site, microtubule stabilization occurs which are being observed in the case of sporozoites. As microtubule dynamicity play crucial role in sporozoite motility and infectivity, our result clearly demonstrated that treatment of sporozoites with these compounds inhibited sporozoite infectivity in hepatocyte cell line HepG2 *in vitro* (**Figure 4a(i)**), as well motility (**Figure 4a(ii)**), suggesting that the sporozoite motility and infectivity depend on dynamicity of the microtubule. We next investigated the effect of these microtubule stabilizers on *P. berghei* gametocyte maturation. Notably, these three compounds: MMV676477, MMV1578136 and MMV1578138 significantly inhibited the gametocyte maturation when tested *ex vivo* **(Figure 4c).** In terms of the mechanism of action of the observed immature gametocytes, it has been suggested that during the final stages of the gametocyte maturation, microtubules depolymerize, resulting in a drop in parasite length and a change in the deformability of the host red blood cell. But, because of the treatment with these compounds, this process was severely affected leading to inhibition of gametocyte maturation. After gametocyte maturation, male gamete undergoes ex-flagellation which releases up to 8 flagellated male gametes from a single male gametocyte. Since, the motile machinery of the sexually active gametes is comprised of flagella that are primarily composed of axonemal microtubules, we next evaluated the inhibitory action of these compounds on male gamete ex-flagellation, and found that they significantly inhibited this process by 70% (3µM), as shown in **Figure 4d.**

Since these compounds inhibited both the maturation and ex-flagellation of gametes, we next looked to determine if they were also capable of inhibiting zygote to ookinete conversion. Interestingly, the compounds: MMV676477, MMV1578136 and MMV1578138 (at 3μM concentration) abolished the maturation of ookinete when tested against *P. berghei ex vivo*. Previously, Taxol and Colchicine MTAs agents have been reported to abolish the transformation of zygotes into ookinetes, however, the inhibitory concentration used was 10μM and 25μM, respectively, which is higher than the current study. Moreover, these compounds also inhibited the sporozoite infectivity and motility, as illustrated in **Figures 4a and 4b**. Their possible mechanism of action at sporozoite stages could be the sub-pellicular microtubule, which undergoes dynamic morphological changes during the invasion and intracellular development. Furthermore, the lack of cytotoxicity displayed by the parent scaffold, MMV676477 against human cell line, coupled with a promising ADME profile (Table 2) support its consideration as a lead compound for further development to ultimately validate *Pf*Tubulin as a therapeutic target.

Overall, this study identified tubulin as a biological target in the malaria parasite which is essential throughout the *Plasmodium* life cycle, and could be targeted to develop multistage anti-malarials. Three MMV compounds: MMV676477, MMV1578136 and MMV1578138 from a pyrazole-pyrimidine series are shown to be of great importance as lead molecules for the development of potential multistage inhibitors to fight against plasmodium drug resistance.

## AUTHOR CONTRIBUTIONS

S. Singh conceived the idea and designed the experiments, wrote and edited the manuscript. GK, RJ, and RKS carried out the experiments, interpreted the results, analysed the experimental data, and wrote the manuscript. MV assisted with the drafting of the text. IK and P assisted in sporozoite isolation. APS directed sporozoite-related investigations. KS assisted with technical recommendations, and JB assisted with chemical procurement and experimental direction.

## ACKNOWLEDGEMENTS

We are thankful to Advanced Instrumentation and Research Facility (AIRF), JNU, New Delhi for Surface Plasmon Resonance (SPR) facility. Funding from National Bioscience Award from the Department of Biotechnology (DBT), Government of India (Sanction No. BT/HRD/NWBA/39/04/2018-19) is acknowledged. GK and RKS are recipients of fellowship from Council of Scientific and Industrial Research (CSIR), Government of India. RJ acknowledges University Grant Commission (UGC) for doctoral research fellowship. The funders had no role in study design, data collection and analysis, decision to publish, or preparation of the manuscript.

